# Thermal flexibility is a repeatable mechanism to cope with environmental stressors in a passerine bird

**DOI:** 10.1101/2021.02.03.429657

**Authors:** Joshua K. Robertson, Gabriela F. Mastromonaco, Gary Burness

## Abstract

1. For many vertebrates, urban environments are characterised by frequent environmental stressors. Coping with such stressors can demand that urban individuals activate energetically costly physiological pathways (e.g. the fight-or-flight response) more regularly than rural-living conspecifics. However, urban environments also commonly demand appreciable expenditure toward thermoregulation, owing to their often extreme climatic variations. To date, whether and how vertebrates can balance expenditure toward both the physiological stress response and thermoregulation, and thus persist in an urbanising world, remains an unanswered and urgent question among ecologists.
2. In some species, changes in body surface temperature (T_s_) and peripheral heat loss (q_Tot_) that accompany the stress response are thought to balance energetic expenditure toward thermoregulation and responding to a stressor. Thus, augmentation of stress-induced thermal responses may be a mechanism by which urban individuals cope with simultaneously high thermoregulatory and stress-physiological demands.
3. We tested whether stress-induced changes in T_s_ and q_Tot_: (1) differed between urban- and rural-origin individuals, (2) reduce thermoregulatory demands in urban individuals relative to rural conspecifics, and (3) meet an essential first criterion for evolutionary responses to selection (variability among, and consistency within, individuals).
4. Using the black-capped chickadee (*Poecile atricapillus*; n = 19), we show that neither rapid nor chronic changes in T_s_ and q_Tot_ following exposure to randomised stressors differed between urban- and rural-origin individuals (n_urban_ = 9; n_rural_ = 10). Nevertheless, we do find that stress-induced changes in T_s_ and q_Tot_ are highly repeatable across chronic time periods (R_Ts_ = 0.61; R_qTot_ = 0.67) and display signatures of stabilising or directional selection (i.e. reduced variability and increase repeatability relative to controls).
5. Our findings suggest that, although urban individuals appear no more able to balance expenditure toward thermoregulation and the stress response than rural conspecifics, the capacity to do so may be subject to selection in some species. To our knowledge this is also the first study to report repeatability of any theorised stress-induced trade-off.

## 1 INTRODUCTION

Over the past 70 years, the global human population has increased by approximately 350% (or approximately 5.1 billion; United Nations 2019). Unlike in previous centuries, the majority of individuals (nearly 54%) now reside in urban environments, and global trends strongly suggest that urban living will increasingly become the norm (reviewed in Lerch 2017). Consequently, land area designated for urban utility is expanding at unprecedented rates and will probably continue to do so over the coming decades (Angel *et al*., 2011). Such expansion cannot, however, occur in a vacuum, and has thus contributed to the widespread reduction in habitat availability and quality for many species (Grimm *et al*. 2008; Seto *et al*. 2012; Freeman *et al*. 2019; lay literature: Thomas 2017). For this reason, understanding whether these species can adapt and persist withinsmodern city-scapes has become a growing priority among modern ecologists and conservationists (e.g. Birnie-Gauvin *et al*. 2016; Ouyang *et al*. 2018).

Yet habitat loss or degradation are not the only challenges faced by species in urban environments. Indeed, urban environments regularly present acute challenges to those residing within, including noise, frequent human interaction, vehicle traffic, and in some cases, elevated depredation and inter- and intra-specific competition (Johnson *et al*. 2012; Hernández-Brito *et al*. 2014; Newsome *et al*. 2015; Vincze *et al*. 2017; reviewed in Lowry *et al*. 2013). Coping with these acute challenges can demand that urban-living individuals activate self-preserving physiological responses (i.e. fight-or-flight responses) more regularly than rural-living conspecifics (Bonier 2012; Watson *et al*. 2017; albeit, often with reduced intensity; Partecke *et al*. 2006; French *et al*. 2008; but see Fokidis *et al*. 2009). While such demands need not inherently translate to a loss of fitness among urban individuals, laboratory studies suggest that their daily metabolic costs are probably raised owing to increased allostatic load (Depke *et al*., 2008; Jimeno *et al*., 2017). In turn, these elevated metabolic demands may enhance susceptibility to wear and tear when resources are restricted or are required to be allocated elsewhere (Romero *et al*., 2009; Breuner & Berk, 2019).

Beyond urban development, many of today’s species face additional and indirect threats associated with a growing human population. effects of anthropogenic climate change on species distribution and trait expression, for example, have now been argued for nearly all taxa (e.g. Barton *et al*. 2016; Mainwaring *et al*. 2017; Pacifici *et al*. 2017; Wan *et al*. 2018), and concerns over the ability of species to adjust to rising and increasingly variable ambient temperatures (Vasseur *et al*., 2014) have been well articulated (e.g. Rutschmann *et al*. 2015; Radchuk *et al*. 2019). In endotherms, increases in both maximal ambient temperature and variability of ambient temperatures can bear notable thermoregulatory costs (Pendlebury *et al*., 2004; du Plessis *et al*., 2012; Smit *et al*., 2018), with those associated with the former being particularly severe in urban environments (Arnfield, 2003). These costs, coupled with expected increases in susceptibility of wear and tear, beg important questions of whether and how endotherms may cope with increasingly urbanised environments in the face of a rapidly changing climate (discussed in Pautasso 2012; Argüeso *et al*. 2015; Brans *et al*. 2017).

To date, several empirical studies have shown that endotherms may adjust their superficial blood-flow, and thus, their body surface temperatures (henceforth, “T_s_”) when exposed to stressors (e.g. Blair *et al*. 1959; Yokoi 1966; Nord & Folkow 2019; Winder *et al*. 2020). In some species, these changes in T_s_ appear to endow individuals with greater heat conservation in the cold, and greater heat dissipation in the warmth, thus reducing their demands for costly thermogenesis or evaporative cooling respectively (Jerem *et al*. 2018; Robertson *et al*. 2020a; Winder *et al*. 2020). In this way, total energetic expenditure may be balanced in challenging environments by allocating energy toward more immediate and higher-cost threats (e.g. the perceived stressors) and away from less immediate and lower-cost threats (e.g. thermal challenges; Jerem *et al*. 2018; Robertson *et al*. 2020a). In urban environments, where individuals regularly contend with both physical and thermal challenges, such flexibility of T_s_ and peripheral heat loss (here, non-evaporative heat-loss; henceforth, “q_Tot_”) could be particularly advantageous, with those capable of enhanced flexibility (particularly during stress exposures) being better able to balance energy expenditure and, therefore, being favoured by selection (see Parsons 2005). Nevertheless, the potential for selection to act on flexibility of T_s_ and q_Tot_ in response to stressors requires that these traits are both variable among individuals, and consistent within individuals (i.e. “repeatable”; reviewed in Boake 1989; Wolak *et al*. 2012). Over the past two decades, numerous studies have reported moderate to high degrees of repeatability among traits associated with the stress response and whole-animal metabolism (Nespolo & Franco 2007; Rensel & Schoech 2011; Müller *et al*. 2018; Boratyński *et al*. 2019; but see Ouyang *et al*. 2011). While these finding strongly suggest that stress-induced changes in T_s_ and q_Tot_ are also likely to be repeatable in endotherms, the degree of this repeatability remains largely unclear (but see Careau *et al*. 2012).

Using the black-capped chickadee (*Poecile atricapillus*, Linnaeus, 1776; henceforth “chickadees”) as a model species, we tested whether flexibility of both T_s_ and q_Tot_ during stress exposure: (1) meet a critical first criterion for responsiveness to selection, and (2) offer opportunities for endotherms to cope with the increased allostatic and thermoregulatory costs of an urbanising environment. More specifically, we hypothesised that stress-induced changes in both T_s_ and q_Tot_: (1) are variable among, and consistent within individuals, (2) provide evidence of current or past selection, and (3) differ between individuals captured from urban and rural environments.

In accordance with our hypotheses, we first predicted that stress-induced changes in both T_s_ and q_Tot_ would be repeatable among individuals. Because thermal responses to stress exposure can be acute (e.g. minutes to hours: Jerem *et al*. 2015; Andreasson *et al*. 2020; Winder *et al*. 2020) or chronic (e.g. days: de Aguiar Bittencourt *et al*. 2015; Herborn *et al*. 2018), and responses across each time-period may provide energetic benefits by enhancing heat dissipation or relaxing costs of thermogenesis (Jerem *et al*. 2018; Herborn *et al*. 2018; Winder *et al*. 2020), we predicted that both acute and chronic changes in T_s_ and q_Tot_ accompanying the stress response would be repeatable among individuals in our sample population. Next, because traits subject to previous or current selection (here, stabilising or directional) are thought to display lower variability and higher repeatability than those that are selectively neutral (e.g. Gibson & Bradley 1974; Lande & Arnold 1983; Boake 1989; Van Homrigh *et al*. 2007; but see Kotiaho *et al*. 2001), we predicted that both T_s_ and q_Tot_ of chickadees would be less variable and more repeatable during stress exposure treatments than during control treatments, after controlling for predictable environmental effects on heat loss (e.g. ambient temperature and relative solar radiation). Finally, because the combined energetic costs associated with the stress response and thermoregulation are expected to be higher in urban environments when compared with rural environments (i.e. in the absence of phenotypic differences between urban and rural individuals; discussed above), we predicted that the magnitude of both acute and chronic changes in T_s_ and q_Tot_ that accompany stress exposures would be larger among urban-origin individuals than rural-origin individuals.

To test our predictions, we exposed chickadees captured from urban and rural environments to both repeated stressors and control conditions across an ambient temperature gradient while monitoring rapid and long-term changes in T_s_ and q_Tot_ by infrared thermography. In small birds, surface tissues at the periorbital region (henceforth “eye region”) are thought to play a critical role in environmental heat exchange (e.g. Hill *et al*. 1980; Powers *et al*. 2015) and both temperature of, and heat loss from this re-gion have previously been shown to respond to stress exposure (e.g. Jerem *et al*. 2015; Ikkatai & Watanabe 2015; Herborn *et al*. 2018; Robertson *et al*. 2020a). We, therefore, chose to use temperature of, and heat loss from, the eye region as our indicators of T_s_ and q_Tot_ in this study.

The capacity of vertebrates to cope with the combined pressures of urbanisation and anthropogenic climate change has been questioned many times (Pautasso, 2012; Argüeso *et al*., 2015; Brans *et al*., 2017). The proximate physiological mechanisms by which vertebrates (here, endotherms) may do so, however, are seldom explored. Ours study, therefore, represents a critical step forward in how ecologists might test the capacity of vertebrates to adapt to an increasingly human-modified world.

## 2 MATERIALS AND METHODS

All methods used for animal capture, sampling, and experimental treatment were approved by the Trent University Animal Care Committee (AUP # 24614) and Environment and Climate Change Canada (permit # 10756E).

### 2.1 Capture, transport, and housing of experimental animals

Chickadees (n = 20; n = 10 females, n = 10 males) used for this experiment were captured within a 100 km^2^ region of south-western Ontario, Canada, between the months of March and April in 2018. To minimise the possibility of kinship between individuals within our sample population, capture efforts were divided across six distinct locations (three urban and three rural), each separated by a minimum distance of 15 km. Urban capture locations included the downtown regions of the cities of Brantford (43.1345°N, 80.3439°W), Cambridge (43.3789°N, 80.3525°W), and Guelph (43.3300°N, 80.1500°W), while rural capture locations included the townships of Corwhin (43.5090°N, 80.0899°W), Erin (43.7617°N, 80.1529°W), and Cayuga (42.9797°N, 79.8745°W; Figure S1). A difference in the mean degree of urbanisation between urban and rural capture locations was validated using methods similar to Thompson et al (2018; see Appendix; Figures S2-S4).

All individuals were captured using modified potter traps (dimensions [L × W × H] = 90 × 70 × 70 cm), baited with sunflower seeds and suet on the day of capture. To further draw individuals to trap locations, we alternately broad-casted chickadee breeding songs and alarm calls from a remote call-box (FoxPro™ Patriot; Lewisville, PA, USA) until at least one individual approached a potter trap by ≤ 4 meters. Upon capture, chickadees were blood sampled (approximately 50 *µ*L) by brachial venipuncture and capillary tube collection, then fitted with one stainless steel, numbered leg ring (size 0) and a unique combination of two, coloured, Darvic leg rings (size 0) for future identification. Each individual was then measured (mass to the nearest 0.1 g using an electronic scale, and, wing cord to the nearest 0.1 mm, left outer tarsus to the nearest 0.1 mm, and head-to-bill to the nearest 0.1 mm using analogue calipers) and secured in a covered flight enclosure (dimensions [L × W × H] = 30 × 30 × 15 cm) for transportation to our long-term housing facility (Ruthven Park National Historic Site, Cayuga, Ontario; ≤ 90 km drive). Blood samples were preserved in a small volume of Queen’s Lysis buffer (500 *µ*L; Seutin *et al*. 1991) for use in genetic sexing (using methods described in Robertson *et al*. 2020a) and were held on ice until storage at 4°C was possible (≤ 2 hours).

Upon arrival to our long-term housing facility, chickadees were haphazardly distributed among four, visually isolated flight enclosures (n = 5 per enclosure; dimensions [L × W × H] = 1.83 × 1.22 × 2.44 m), each equipped with one white cedar tree (*Thuja occidentalis*), two perching branches (raised to approximately 1.50 and 1.80 m above ground) and a raised feeding platform (400 cm^2^) at which food was provided *ad libitum* through an opaque hinged door for the duration of the experiment (Figure 1). Food provided included sunflower seed, safflower seed, shelled peanuts, boiled egg, apple pieces, house crickets (*Acheta domesticus*), meals worms (*Tenebrio molitor*) and Mazuri (St Louis, MO, USA) Small Bird Maintenance diet. Water was also provided *ad libitum* across our experiment through opaque hinged doors. All individuals were given a minimum of 2 weeks to acclimate to enclosures and social groups prior to the onset of experimentation.

**FIGURE 1.**
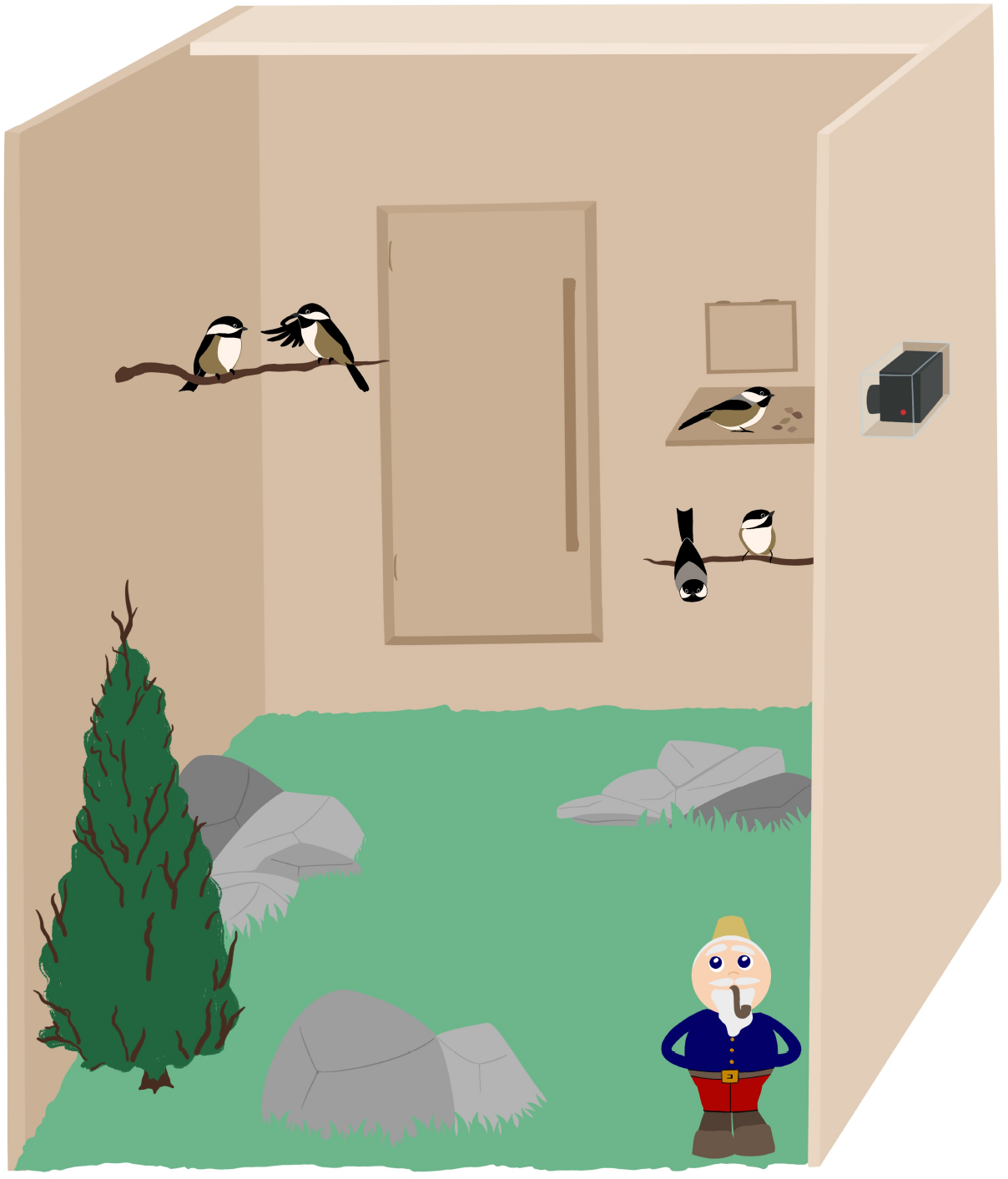
Depiction of experimental stress exposure (novel object) and infrared thermographic imaging in a selected flight enclosure. Black-capped chickadees (n = 5) within a given flight enclosure were simultaneously exposed to each individual stressor (here, the presence of a garden gnome), while individuals at raised feeding platforms were passively imaged with a remotely activated infrared thermographic camera.

The minimum and maximum ambient temperatures observed during our study were 3.0 °C and 38.5°C respectively, and day-length (duration between civil dawn and civil dusk) ranged from approximately 14.75 hours to 16.5 hours.

### 2.2 Experimental stress exposure

To test repeatability of stress-induced thermal responses within and among individuals, we used a paired experimental design wherein each individual was exposed to both a thirty-day control treatment and a thirty-day stress exposure treatment, with treatments separated by an additional two-day control period (total experimental duration = 62 days). To control for possible effects of treatment order on stress-induced thermal responses, half of our sample population (n = 10 across two flight enclosures) was exposed to control treatments followed by stress-exposure treatments, while the second half of our sample population (n = 10 across two flight enclosures) was concurrently exposed to a reversed treatment order (i.e. stress-exposure treatments followed by control treatments).

Each day, individuals within stress exposure treatments were exposed to 5 or 6 experimental stressors, with each being applied for 20 minutes and being separated from previous and subsequent stress exposures by ≥ 1 hour (similar to Rich & Romero 2005). Timing and type of experimental stressors were randomly selected each day to minimise the potential for habituation to each given stressor type. Experimental stressors included the presence of a novel object (a garden gnome), presence of a mock predator (a taxidermically mounted Cooper’s hawk; *Accipiter cooperii*), capture and restraint in an opaque fabric bag, presence of a human, covering of a given flight enclosure with an opaque fabric (simulating extreme, inclement weather), and presence of a taxidermically mounted conspecific fixed to the feeding platform of a given flight enclosure (simulating a novel, dominant individual). In a previous study, chickadees exposed to our randomised stressor protocol displayed a significant reduction in feeding rate and mass, and regularly evoked alarm calls (Robertson *et al*. 2020b), providing strong support for protocol efficacy. Endocrine responses to stressor types were not measured to circumvent effects of blood sampling on surface temperature measurements and stress perception among sampled individuals. Individuals exposed to control treatments were left undisturbed in an adjacent flight enclosure and blind to experimenter presence.

Because flight enclosures were not auditorily segregated, estimated thermal responses to stress exposure in this study (i.e., the interaction between time or ambient temperature and treatment type) are expected to be conservative.

### 2.3 Infrared thermography, body surface temperature estimation, and heat transfer estimation

We monitored T_s_ and q_Tot_ of chickadees indirectly using remote infra-red thermography (thermographic camera: FLIR VueProR™, 13 mm, 226 × 356 resolution: accuracy = *±* 5%; image frequency = 1 Hz). Specifically, we captured infrared thermographic images (radiometric JPEGs) of individuals at feeding platforms across the duration of our experiment from weather-proofed camera boxes mounted to the exterior of enclosure walls (0.5 m distance). To minimise temporal bias of thermographic imaging among social groups, we rotated our thermographic camera cardinally clockwise among flight enclosures each day, with filming durations persisting for approximately one hour per enclosure, and the first flight enclosure to receive thermographic filming being rotated each day. Because leg-ring combinations could not be readily distinguished from thermographic images, we also captured digital video (camera: Action Cam™, Sony, Toronto, Ontario, CA) of feeding individuals in parallel to themographic images to permit individual identification. All thermographic imaging and digital video used in this study were captured between 08:00h and 16:00h of each day.

Estimation of an object’s T_s_, and consequently rate of heat transfer (q_Tot_) by infrared thermography requires that local ambient temperature and relative humidity are known (Minkina & Dudzik, 2009; Tattersall, 2016). We therefore monitored ambient temperature at enclosures subjected to thermographic filming using a ThermoChron iButton™ (Maxim Integrated, DS1922L-F5, San Jose, CA, USA) placed in the shade, at a frequency of 1 reading/5 minutes. Relative humidity readings were collected from a nearby weather station operated by Environment and Climate Change Canada (station identity = Hamilton A, 22 km from the experimental holding location) at the maximum available frequency of 1 reading/hour.

To estimate T_s_ from infrared thermographic images, we followed methods described elsewhere (Robertson *et al*., 2020a). Specifically, raw infra-red radiance (kW/m^2^) values per pixel were manually extracted in R statistical software (version 3.6.1; R Core Team 2019) then first converted to temperature (°C) per pixel according to Planck’s law, ambient temperature, and humidity estimates at the time of image capture, and equations outlined elsewhere (Minkina & Dudzik 2009; Tattersall 2016). Emissivity of the eye region of chickadees was assumed to be fixed at 0.95 according to estimates made for integument of Canadian and snow geese (*Branta canadensis* and *Chen caerulescens* respectively; Best & Fowler 1981). Following their estimation, temperature values per pixel were then integrated into FITS matrices using the R package FIT-Sio (version 2.1.0; Harris 2016; one matrix per thermographic image), and eye region T_s_ values (here, maximum temperature values, as per Jerem *et al*. 2015) were manually extracted from within matrices using the open-sourced software FIJI (Schindelin *et al*. 2012; average size of eye region *≈* 230 pixels). To minimise underestimation of T_s_ as a consequence of image blurring, only values extracted from individuals that were stationary during image capture were included in our final data (Tattersall, 2016). Although recent studies have shown that the rotation of an object within an infra-red thermographic image may influence estimates of its surface temperature (PlayàMontmany & Tattersall, 2021), rotation of chickadees at feeding platforms was unlikely to differ systematically between our control and treatment groups and was therefore not estimated in this study.

To estimate q_Tot_ (mW) from T_s_ measurements, we followed equations described by McCafferty *et al*. (2011) and Nord & Nilsson (2019). Here, however, values for the kinematic viscosity of air (m^2^/S; at an assumed atmospheric pressure of 101.325 kPa) and the thermal expansion coefficient of air (1/K) were estimated for each given ambient temperature using the R packages “bigleaf” and “Thermimage” respectively (Knauer *et al*., 2018; Tattersall, 2019). For this study, q_Tot_ was assumed to equal the sum of convective and radiative heat transfer, owing to both the minimal effects of wind-speed in our flight enclosures, and low likelihood of heat transfer between the eye region and any medium other than air during our experiment. Surface area of the eye region was estimated as 0.864 cm^2^ (ovoid with horizontal diameter of 1.1 cm and vertical diameter of 1.0 cm), and contours within the eye region were considered negligible. Final q_Tot_ estimates were multiplied by two to represent q_Tot_ across both eye regions.

### 2.4 Statistical analyses

All statistical analyses were conducted in R software (version 3.6.1; R Core Team 2019 with each generalised additive mixed-effects model (“GAMM”) constructed in the package “brms” (version 2.13.3; Bürkner 2017). Additionally, all models were run using Markov Chain Monte Carlo (MCMC) sampling, with 4 Markov chains, 10000 chain iterations, and 1000 warm-up iterations to maximise mixing and convergence of Markov chains. Final iterations were thinned by 10 to account for possible auto-correlation between MCMC draws, and models were validated by visually diagnosing residual distributions and trace plots. 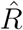 values for all model parameters fell between 0.99 and 1.01, and the ratio of effective sample sizes to our total sample size were greater than 0.65 for each parameter. Lastly, all figures were produced in R using the package “ggplot2” (version 3.3.2; Wickham 2016), and one individual (a female captured in an urban environment) was removed due to an unusually small sample size (n = 19 thermographic images).

#### 2.4.1 Thermal responses to stress exposure among individuals

To first test whether acute and chronic changes in T_s_ accompanying the stress responses were repeatable among individuals, we constructed two Bayesian hierarchical GAMMs wherein we estimated both global responses and individual-level responses to stress exposure across acute and chronic time scales. In both models, temperature of the eye region of individuals (°C; Gaussian distributed) was included as the response variable, and treatment type (i.e. stress exposure or control) and sex were included as linear, population-level predictors to account for the influence of each on eye region temperature measurements. Additionally, flight enclosure identity, date of thermographic image capture, and individual identity were included in each model as group-level intercepts to account for statistical non-independence between measurements collected from the same flight enclosure, day, and individual, and a group-level slope for time of day per flight enclosure orientation (i.e. east facing or west facing) was included to account for differential exposure to solar radiation within east- and west-facing enclosures across time.

In our model predicting acute thermal responses to stress-exposure, time post stress exposure (seconds), ambient temperature, and time of day (hour) were each included as population level predictors. Because acute, stress-induced changes in T_s_ at the eye region are thought to be non-linear (Jerem *et al*., 2015, 2019), time post-stress exposure was included as a cyclic cubic regression spline with 5 knots fixed at −1200, 0, 1200, 2400, and 3600 seconds to evenly distribute model fitting across each phase of stress exposure (i.e. before, during, and after exposure). Here, a cyclic regression spline was chosen to capture expected returns to baseline T_s_ (as reported for blue tits, *Cyanistes caeruleus*; Jerem *et al*. 2019) following 40 minute recovery periods. To permit comparisons between stress exposed and control treatments, we paired enclosures such that time post stress exposure for an enclosure experiencing a control treatment was considered to be equivalent to that of the nearest enclosure experiencing a stress exposure treatment and equivalent cardinal orientation (i.e. west- or east-facing). As such, our comparisons between treatments account for indirect effects of experimental stress exposures on nearby control individuals.

In endotherms, T_s_ is expected to display non-linear relationships with both ambient temperature and time of day owing to peripheral thermoregulatory processes (i.e. cold-induced vasoconstriction and warm-induced vasodilation) and circadian rhythms (Richards, 1971; Cooper & Gessaman, 2005) respectively. Ambient temperature and time of day were therefore included as natural cubic and thinplate regression splines respectively, each with 4 knots to minimise risk of model over-fitting. Knots for our ambient temperature spline were evenly spaced by quantiles to uniformly capture trends in eye region temperature at ambient temperatures below, within, and above thermoneutrality for our study species (Grossman & West, 1977). Because we did not have *a priori* assumptions for knot positions for our time of day spline, knot positions were chosen by truncated eigen decomposition (Wood, 2003). To control for differential effects of treatment type on T_s_ across time (Jerem *et al*., 2015, 2019) and ambient temperature (Robertson *et al*. 2020a), population-level interactions between treatment type and ambient temperature, and treatment type and post stress exposure were also included as model predictors, along with an interaction between treatment type and the tensor product (*⊗*) between ambient temperature and time post stress exposure to account for the influence of ambient temperature on acute thermal responses to stress exposure at the skin (Nord & Folkow, 2019). All interaction terms were penalised on the first derivative to minimise the potential for concurvity between interaction terms and main effects. Finally, to estimate differences in acute, stress-induced changes in T_s_ among individuals, group-level slopes for time post stress exposure and the interaction between time post-stress exposure and treatment type were included for each individual. Correlations between adjacent T_s_ measurements was corrected using a type-I autoregressive (AR1) correlation structure with an estimated rho (*ρ*) of 0.69, and residual error was estimated independently for each treatment type.

In our model predicting chronic stress-induced changes in T_s_, group-level predictors remained as described above but with minor adjustments. Specifically, all predictors including time post stress exposure (i.e. as a main effect or interactive effective) were excluded from our model to permit assessment of long-term, but not short-term trends in T_s_ according to treatment type. Furthermore, to estimate differences in chronic stress-induced changes in T_s_ among individuals, group-level slopes for ambient temperature and the interaction between ambient temperature and treatment type was included per individual. Here, ambient temperature was mean-centered and scaled to 2 times the standard deviation (as per Araya-Ajoy *et al*. 2015) to allow for individual slopes to be estimated with respect to our average environmental conditions. Again, correlations between adjacent T_s_ measurements was corrected using an AR1 correlation structure (*ρ* = 0.69), and residual error was estimated separately per treatment.

Because rates of peripheral heat transfer (q_Tot_) are proportional to T_s_ at given ambient temperatures, both acute and chronic changes in q_Tot_ accompanying stress exposure treatments were modeled as described above. In these models, however, q_Tot_ was used as the response variable (mW; Gaussian distributed) in place of T_s_.

In all hierarchical models, we used informed priors for our population intercept, our coefficients for treatment (linear), sex (linear), ambient temperature (first order, linear), and our values for spline smoothness (*ϕ*), with prior distributions being informed by another study using black-capped chickadees (Robertson *et al*., 2020a). For our model intercepts, we used gamma distributed priors with *α* values of 60 and 50 (T_s_ models and q_Tot_ models respectively), and *β* values of 2 (both T_s_ models and q_Tot_ models) thus assuming positive T_s_ and q_Tot_ values at an ambient temperature of 0°C, with peak densities of approximately 30°C and 25 mW respectively. In all models, priors for treatment type and sex were normally distributed with means of 0 and −1 respectively, and standard deviations of 2.5, while those for *ϕ* were gamma distributed with *α* = 2, and *β* = 0.5 owing to low expected “wiggliness” in our smooth terms. Lastly, for our first order slope of ambient temperature, we used gamma distributed priors (*α* = 4, *β* = 2) in our models pertaining to T_s_ and normally distributed priors (mean = −5, s.d. = 5) in our models pertaining to q_Tot_ because the relationship between ambient temperature and T_s_ is expected to be positive, while that between ambient temperature and q_Tot_ is expected to be negative. Uninformative priors were used for all other model parameters; specifically, priors for the standard deviation of population level and group level predictors followed student’s t distributions with 3 degree of freedom, location parameters of 0 and a scale factors of 3.4. Similarly, priors for sigma parameters also followed student’s t distributions with 3 degrees of freedom and location parameters of 0, however, scale factors were reduced to 2.5.

#### 2.4.2 Repeatability estimates

To calculate repeatability of stress-induced changes in T_s_ and q_Tot_, we followed methods described by Araya-Ajoy *et al*. (2015). Their methods, however, are largely descriptive and do not test the presence or absence of trait repeatability within an experimental context. To correct for this, we constructed null models (i.e. models with individual identities scrambled) for T_s_ and q_Tot_ across both acute and chronic time-periods, then compared mean repeatability estimates (per Markov chain iteration) acquired from true and null model posterior distributions. Here, a significant increase in repeatability values derived from true models relative to those derived from null models suggests that true repeatability values could not be explained by biases in the experimental process alone. Null models were constructed by randomly allocating individual identities to each T_s_ and q_Tot_ estimate, then re-running hierarchical models as described above (Figure 2). To control for possible effects of treatment order during identity randomisation, we limited possible identity assignments to individuals that had experienced the same treatment order as the true individual from which the T_s_ or q_Tot_ values were obtained. Mean repeatability estimates were then compared between our true and null models using two, one-way, non-linear hypothesis tests in the R package “brms” (Bürkner, 2017). For all hypothesis tests, priors for true and null repeatability estimates were beta distributed with peaks at 0 (*α* = 1, and *β* = 4). Bayes factors (K), representing support for true repeatability estimates being greater than null repeatability estimates, were calculated from each hypothesis test using the Savage-Dickey density ratio method (Wagenmakers *et al*., 2010).

**FIGURE 2.**
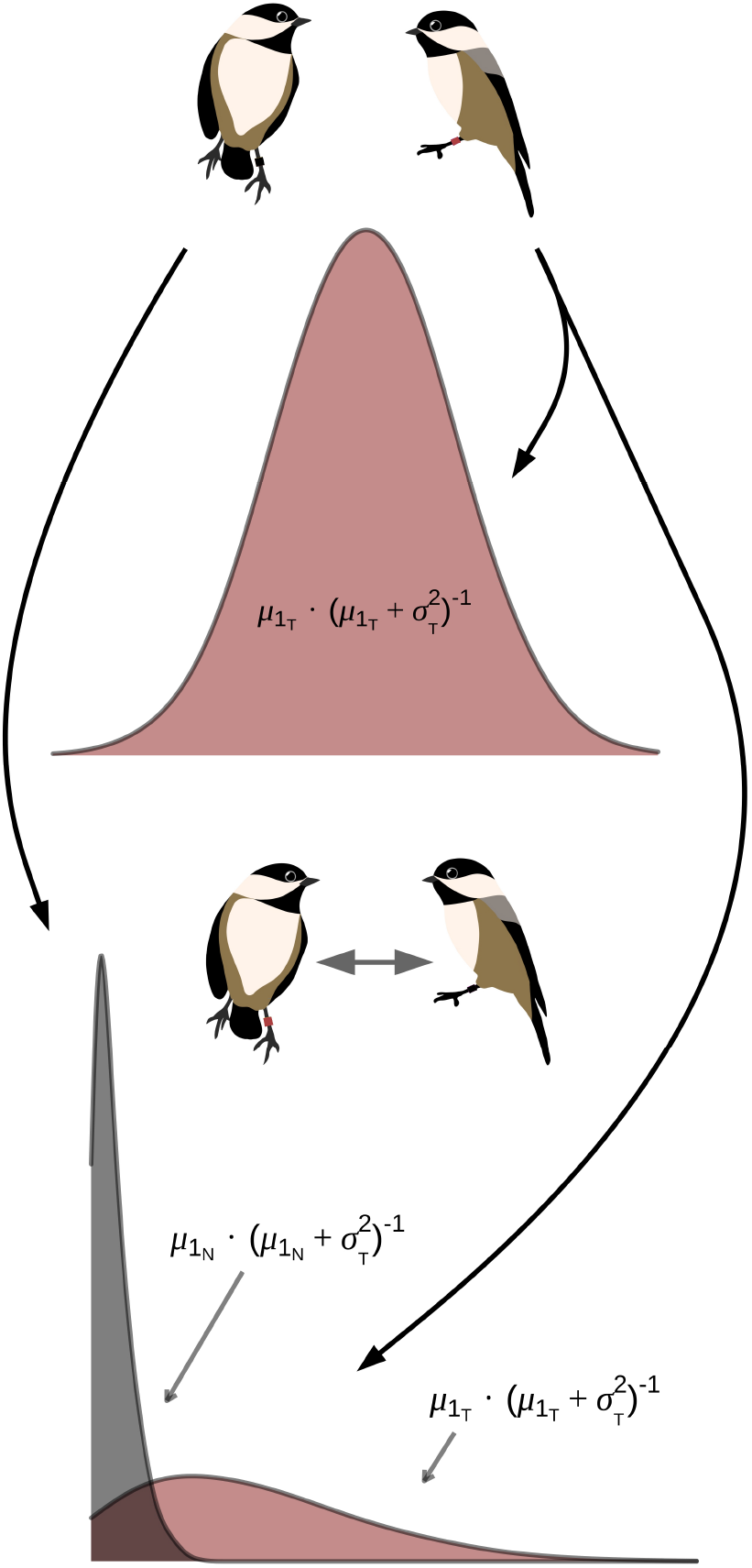
Method used to test for repeatability of stress-induced thermal responses among black-capped chickadees, while controlling for possible biases in the experimental process. Repeatability values were calculated from a true model (maroon; subscripted “T”) using methods described by Araya-Ajoy et al (2015). Individual identities were then scrambled to produce a null model (grey; subscripted “N”), from which repeatability values were again calculated as described above. Final repeatability estimates from true and null models were compared statistically.

#### 2.4.3 Effects of stress exposure on repeatability estimates

Traits under stabilising or directional selection are thought to display lower variability than those that are selectively neutral (e.g. Gibson & Bradley 1974; Lande & Arnold 1983; Van Homrigh *et al*. 2007; but see Kotiaho *et al*. 2001). Furthermore, the potential for traits to respond to selection is contingent upon trait expression being consistent across time (e.g. repeatable; Dochtermann *et al*. 2015; but see Dohm 2002). Thus, the presence of both high repeatability (R) and relatively low residual variation (“*ϵ*” in a linear or additive model) is suggestive of previous or current selection acting upon a trait’s expression, if all other environmental variables and sources of measurement error are controlled (i.e wherein *ϵ* is the sum of residual variation explained by external environmental factors, measurement error, and within-individual variability; suggestive in Gibson & Bradley 1974 and Boake 1989). In our experiment, both stress-exposed and control individuals experienced the same environmental conditions, and measurement error around T_s_ and q_Tot_ was unlikely to differ systemically between stress-exposed and control treatments. Thus, to test for evidence of enhanced stabilising or directional selection (past or current) on the expression of T_s_ and q_Tot_ during stress exposure relative to resting conditions, we compared error and repeatability estimates obtained for stress-exposed and control treatments across both short and long-term time-frames (e.g. acute and chronic, respectively). To do so, both error and repeatability estimates drawn from posterior distributions of acute and chronic models (pertaining to both T_s_ and q_Tot_; described above) were compared using one-way, non-linear hypothesis tests as described previously (subsection “Repeatability estimates”). Priors for repeatability and error estimates under control and stress-exposed conditions were beta (*α* = 1; *β* = 4) and normally distributed (mean = 0, s.d. = 0.25) respectively. Again, Bayes factors were calculated for each test using the Savage-Dickey density ratio method (Wagenmakers *et al*., 2010), with results representing relative support for either decreased error or increased repeatability within stress exposure treatments when compared with control treatments.

#### 2.4.4 Effects of urbanisation on stress-induced thermal responses

To test whether flexible changes in T_s_ and q_Tot_ accompanying acute stress exposures differed between urban and rural chickadees, we first extracted mean coefficients for the interactions between treatment type and time post stress exposure for each individual from the posterior distributions of our acute models. Mean coefficients were then compared between capture ecotypes using Bayesian “ANOVAs” in the R package “BayesFactor” (version 0.9.12.4.2; Morey *et al*. 2019) with capture location (one of six) included as a group-level intercept. To test whether chronic changes in T_s_ and q_Tot_ following stress exposures differed between individuals from urban and rural locations, we used a similar approach, however, mean coefficients for the interactions between ambient temperature and treatment type were extracted from posterior distributions and used as response values. Priors for the effect of capture ecotype and capture location on individual slopes were weak and Cauchy distributed with scale parameters of 2^1/2^ and 1 respectively, while Jeffreys priors were used for our intercept and residual error term (*τ*) (Rouder *et al*., 2012).

## 3 RESULTS

Credible intervals (95%) are reported for model coefficients in crotchets. All reported means are marginal and are given *±* one standard deviation (s.d.).

### 3.1 Stress-induced changes in body surface temperature and peripheral heat loss are repeatable

Our analyses detected rapid and pronounced changes in both eye region temperature (T_s_) and heat loss from the eye region (q_Tot_) of chickadees following stress exposure (T_s_: *β* = 1.68 [0.36, 4.58]; q_Tot_: *β* = 2.79 [0.86, 6.90]; Table 1). Similar and simultaneous changes in T_s_ and q_Tot_ were not detected in nearby control individuals (Table 1). Interestingly, the magnitude and direction of stress-induced T_s_ and q_Tot_ responses were dependent upon ambient temperature (T_s_: *ϕ* = 4.85 [0.62, 10.90]; q_Tot_: *ϕ* = 6.58 [0.64, 15.50]; Table 1). Specifically, at low ambient temperatures (i.e. those below thermoneutrality; < 14°C), individuals exposed to stressors displayed rapid and transient increases in T_s_ and q_Tot_, with elevations in T_s_ and q_Tot_ persisting for approximately 30 minutes (1800 seconds) after stressor completion (Figures 3a and 3b). At out lowest observed ambient temperature (3°C), T_s_ among stress-exposed individuals increased by an average of 5.53°C *±* 0.154°C (with respect to baseline measurements) immediately upon stressor completion (Figure 3a), and this increase corresponded to a rise in q_Tot_ of 11.50 *±* 0.24 mW (Figure 3b). In contrast, at high ambient temperatures (i.e. those above thermoneutrality; > 30°C), an inverted response among stress exposed individuals was detected, with individuals displaying rapid and transient reductions in T_s_ and q_Tot_ (Figures 3a and 3b) in response to stress exposures (albeit small). At these ambient temperatures, decreases in T_s_ and q_Tot_ persisted for approximately 20 minutes (1200 seconds) following stressor completion, with mean T_s_ and q_Tot_ decreasing by approximately 1.15°C *±* 0.152°C and 2.23 *±* 0.24 mW respectively at our highest observed ambient temperature (38.5°C; again, with respect to baseline measurements) upon stressor completion (Figures 3a and 3b). A small effect of time post stress exposure on both T_s_ and q_Tot_ among control individuals was detected (*ϕ* = 0.45 [0.03, 1.90]; Table 1), however, neither increases nor decreases in T_s_ and q_Tot_ were detectable following onset of stress exposures (here, in the nearest-by flight enclosures designated for stress exposure treatments) above or below the thermoneutal zone (Figure 3b). Neither T_s_ nor q_Tot_ differed between sexes (T_s_: *β*_Sex_ = −0.08 [-0.46, 0.33]; q_Tot_: *β*_Sex_ = −0.17 [-0.86, 0.49]; Table 1), and treatment type alone did not influence each value (T_s_:*β*_Treatment_ = 0.26 [-0.19, 0.99]; q_Tot_: *β*_Treatment_ = 0.29 [-0.28, 1.25]; Table 1).

**TABLE 1.** Acute effects of stress exposure on eye region temperature (T_s_) and dry heat transfer (q_Tot_) of black-capped chickadees (n = 19; n = 9 females, n = 10 males); results of two hierarchical GAMMs. Obelisks (†) represent smooth terms, for which estimates refer to the degree of smoothness (*ϕ*: 0 = linear slope). Estimates for remaining population-level terms represent linear slopes, while those for group-level effects represent standard deviations. Degree of smoothness and 95% credible intervals (“CIs”) for tensor products represent means across penalisation groupings, and effective sample sizes represent sums across groupings. Eye region temperature measurements were estimated from infrared thermographic images (n = 5599) captured across 60 days. T_s_ model: R^2^ = 0.85; q_Tot_ model: R^2^ = 0.94. Asterisks (*) represent statistically significant terms (95% credible intervals do not cross zero).

| Population-level Predictors |  |  |  |
| --- | --- | --- | --- |
| Term | T<sub>s</sub> Estimate<br>[95% CIs] | q<sub>Tot</sub> Estimate<br>[95% CIs] | Effective Sample Size<br>(T<sub>s</sub>/q<sub>Tot</sub>) |
| Intercept* | 33.09 [30.84, 35.10] | 19.02 [15.16, 25.30] | 3644/3600 |
| Treatment | 0.26 [-0.19, 0.99] | 0.29 [-0.28, 1.25] | 3726/3917 |
| Sex (Male) | -0.08 [-0.46, 0.33] | -0.17 [-0.86, 0.49] | 3536/3714 |
| <sup>†</sup>Ambient<br>Temperature* | 1.63 [0.38, 5.03] | 1.38 [0.16, 5.00] | 3537/3680 |
| <sup>†</sup>Ambient<br>Temperature:<br>Treatment* | 1.47 [0.19, 5.07] | 1.87 [0.29, 5.74] | 2870/3191 |
| <sup>†</sup>Time Post Stress<br>Exposure* | 0.45 [0.03, 1.90] | 0.65 [0.05, 2.52] | 3273/3440 |
| <sup>†</sup>Time Post Stress<br>Exposure: Treatment* | 1.68 [0.36, 4.58] | 2.79 [0.86, 6.90] | 3566/3679 |
| <sup>†</sup>[Time Post Stress<br>Exposure $\otimes$ Ambient<br>Temperature]:<br>Treatment* | 4.85 [0.62, 10.90] | 6.58 [0.64, 15.50] | 10141/10674 |
| <sup>†</sup>Hour $\otimes$ Orientation* | 3.68 [0.65, 10.10] | 4.19 [0.64, 10.70] | 10571/10674 |
| Group-level Predictors |  |  |  |
| Bird Identity | 0.32 [0.20, 0.50] | 0.56 [0.35, 0.87] | 3763/3121 |
| Date of Photo | 1.79 [1.37, 2.32] | 3.30 [2.50, 4.20] | 3254/3101 |
| Flight Enclosure<br>Identity | 1.51 [0.34, 4.19] | 5.06 [0.96, 14.19] | 3658/3486 |
| Bird Identity: Time<br>Post Stress Exposure<br>(Control) | 0.36 [0.11, 0.56] | 0.49 [0.14, 0.90] | 3397/3496 |
| Bird Identity: Time<br>Post Stress Exposure<br>(Stress Exposed) | 0.46 [0.21, 0.81] | 0.71 [0.28, 1.26] | 3459/3390 |
| Residual Variance and Repeatability |  |  |  |
| $\sigma_{\text{Control}}$ | 1.21 [1.19, 1.24] | 2.09 [2.06, 2.13] | 3420/3917 |
| $\sigma_{\text{Stress exposure}}$ | 1.18 [1.14, 1.22] | 2.06 [1.99, 2.12] | 3542/3679 |
| R<sub>Control</sub> | 0.07 [0.01, 0.18] | 0.06 [0.01, 0.16] | 3420/3917 |
| R<sub>Stress exposure</sub> | 0.14 [0.03, 0.32] | 0.11 [0.02, 0.27] | 3542/3679 |

**FIGURE 3.**
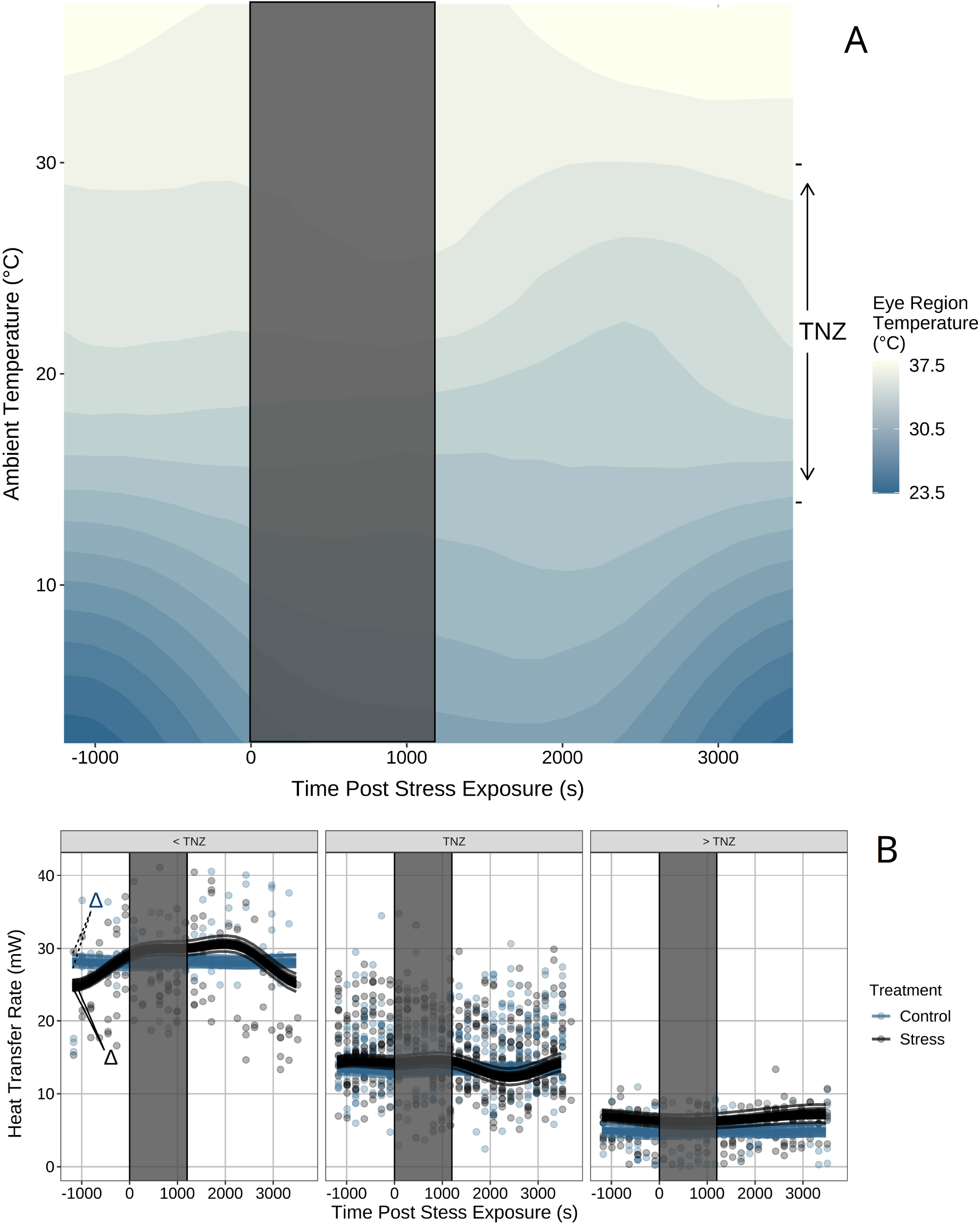
Acute changes in eye region temperature (T_s_) and dry heat transfer (q_Tot_) following stress exposure in black-capped chickadee (n = 19) across ambient temperature. **A** | Average change in T_s_ following stress exposure across ambient temperature (°C) and time since exposure (s). Averages are derived from a Bayesian generalised additive mixed effects model (GAMM) and are marginalised across all other model predictors. T_s_ decreases after stress exposure at ambient temperatures below thermoneutrality, and increases after stress exposure at ambient temperatures above thermoneutrality. **B** | Changes in q_Tot_ of black-capped chickadees across both control and stress-exposed treatments, where slopes per treatment are permitted to vary among individuals. Each line represents the trend for a given individual at temperatures below, within, and above the thermoneutral zone (TNZ; estimated from Grossman and West, 1977), as predicted from a Bayesian GAMM. Dots represent averages per individual across 3 minutes of observation. Both trend lines and dots represent averages for each ambient temperature grouping (< TNZ, TNZ, > TNZ). Grey rectangles in panels A and B represent time when stress exposure treatments were applied in stress-exposed treatment groups. Bold black lines (solid and dashed) and accompanying delta (*δ*) symbols indicate the spread of correlations between time post stress exposure and q_Tot_ across individuals, in control and stress exposure treatments respectively. T_s_ and q_Tot_ were estimated by infra-red thermography (n = 5832 images) across 60 days.

Beyond the acute responses, our analyses also detected chronic effects of stress exposures on T_s_ and q_Tot_ across our sample population (T_s_ model: *β* = 1.81, [0.32, 5.58]; q_Tot_ model: *β* = 2.51, [0.61, 6.92]; Table 2). Specifically, both T_s_ and q_Tot_ of stressexposed individuals decreased at low ambient temperatures and increased at high ambient temperatures relative to controls (Table 2; Figure 4). On average, T_s_ was 1.89°C *±* 1.22°C lower in stress-exposed individuals than control individuals at our lowest observed ambient temperature, and 1.64°C *±*0.95°C higher in stress-exposed individuals than control individuals at our highest observed ambient temperature. Such trends in T_s_ corresponded to reductions in q_Tot_ of approximately 3.75*±* 2.56 mW at our lowest observed ambient temperature, and increases in q_Tot_ of approximately 2.56 *±* 1.99 mW at out highest observed ambient temperature among stress exposed individuals relative to controls (Figure 4). Similar to our results pertaining to acute thermal responses, neither T_s_ nor q_Tot_ differed between sexes in our chronic model (T_s_: *β*_Sex_ = 0.02 [-0.41, 0.44]; q_Tot_ model: *β*_Sex_ = 0.03 [-0.71, 0.76]; Table 2) and no effect of treatment alone on T_s_ or q_Tot_ was detected (T_s_: *β*_Treatment_ = 0.02 [-0.16, 0.20]; q_Tot_: *β*_Treatment_ = 0.00 [-0.29, 0.29]).

**TABLE 2.** Chronic effects of stress exposure on eye region temperature (T_s_) and dry heat transfer (q_Tot_) of black-capped chickadees across ambient temperature (n = 19; n = 9 females, n = 10 males); results of a hierarchical, Bayesian GAMMs. Obelisks (*†*) represent smooth terms, for which estimates refer to the degree of smoothness (*ϕ*: 0 = linear slope). Estimates for remaining population-level terms represent linear slopes, while those for group-level effects represent standard deviation explained by respective terms. Again, degree of smoothness and 95% credible intervals (“CIs”) for tensor products represent means across penalisation groupings, and effective sample sizes represent sums across groupings. Eye region temperature measurements were estimated from infrared thermographic images (n = 5832) captured across 60 days. T_s_ model: R^2^ = 0.85; q_Tot_ model: R^2^ = 0.94. Asterisks (*) represent statistically significant terms (95% credible intervals do not cross zero).

| Population-level Predictors |  |  |  |
| --- | --- | --- | --- |
| Term | T<sub>s</sub> Estimate<br>[95% CIs] | q<sub>Tot</sub><br>[95% CIs] | Effective Sample Size<br>(T<sub>s</sub>/q<sub>Tot</sub>) |
| Intercept* | 32.90 [30.73, 34.75] | 18.68 [14.87, 25.23] | 3479/3419 |
| Treatment | 0.02 [-0.16, 0.20] | 0.00 [-0.29, 0.29] | 3628/3370 |
| Sex (Male) | 0.02 [-0.41, 0.44] | 0.03 [-0.71, 0.76] | 3387/3628 |
| <sup>†</sup>Ambient<br>Temperature* | 1.57 [0.31, 5.31] | 1.28 [0.12, 4.76] | 3742/3425 |
| <sup>†</sup>Ambient<br>Temperature:<br>Treatment* | 1.81 [0.32, 5.58] | 2.51 [0.61, 6.92] | 3608/3299 |
| <sup>†</sup>Hour $\otimes$ Orientation* | 3.41 [0.72, 8.48] | 4.17 [0.81, 9.51] | 7206/6862 |
| Group-level Predictors |  |  |  |
| Bird Identity | 0.35 [0.23, 0.57] | 0.62 [0.40, 1.00] | 3551/3263 |
| Date of Photo | 1.83 [1.40, 2.36] | 3.31 [2.56, 4.26] | 3598/3470 |
| Flight Enclosure<br>Identity | 1.50 [0.31, 4.36] | 4.85 [0.81, 13.86] | 3467/3508 |
| Bird Identity:<br>Ambient Temperature<br>(Control) | 0.88 [0.54, 1.36] | 1.76 [1.09, 2.74] | 3633/3507 |
| Bird Identity:<br>Ambient Temperature<br>(Stress exposure) | 1.52 [0.85, 2.42] | 3.05 [1.82, 4.74] | 3608/3458 |
| Residual Variance and Repeatability |  |  |  |
| $\sigma_{\text{Control}}$ | 1.20 [1.18, 1.23] | 2.07 [2.04, 2.11] | 3503/3846 |
| $\sigma_{\text{Stress exposure}}$ | 1.17 [1.13, 1.21] | 2.04 [1.98, 2.10] | 3297/3461 |
| R<sub>Control</sub> | 0.34 [0.17, 0.56] | 0.41 [0.22, 0.63] | 3503/3846 |
| R<sub>Stress exposure</sub> | 0.61 [0.35, 0.81] | 0.67 [0.44, 0.84] | 3297/3461 |

**FIGURE 4.**
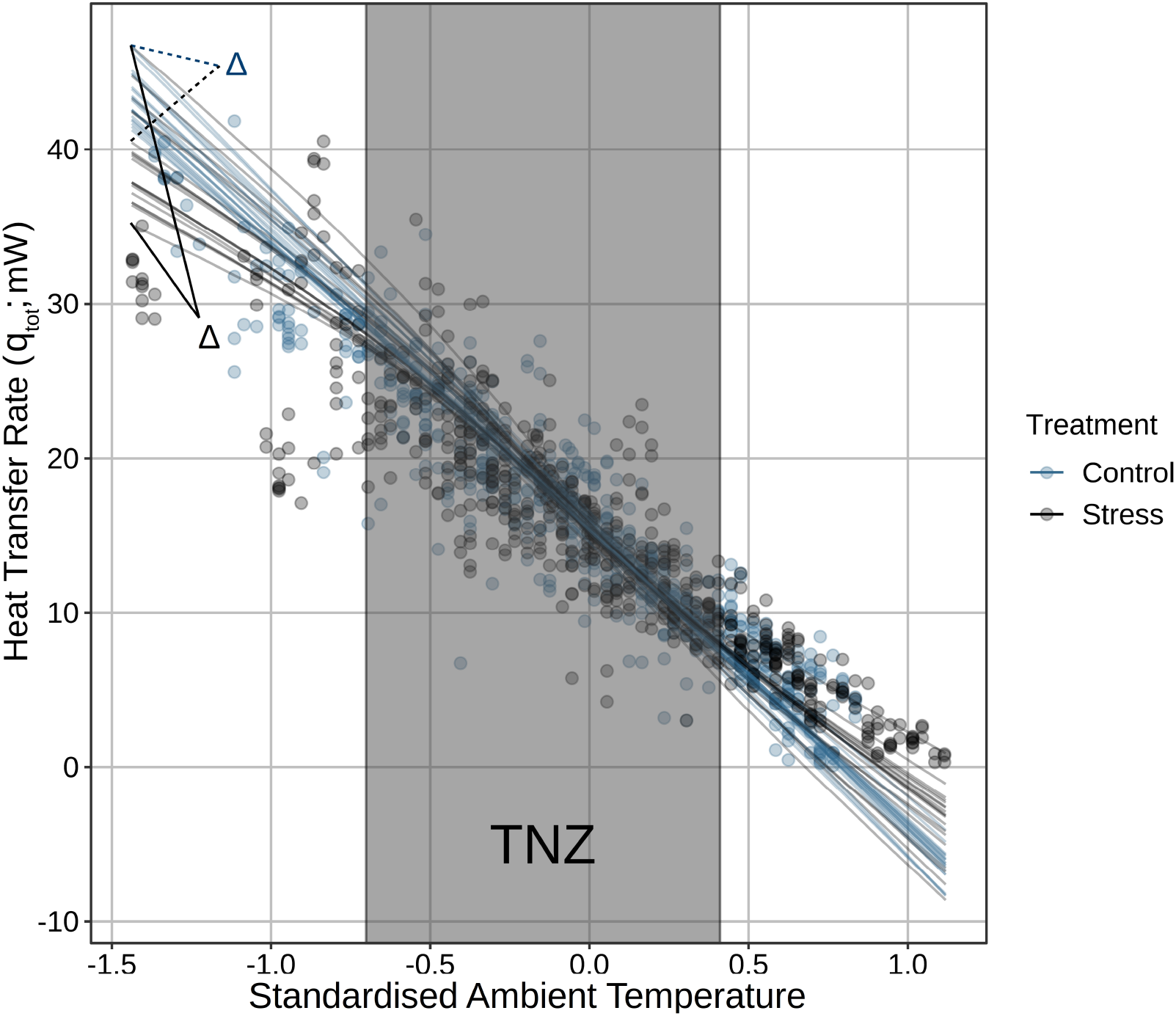
Chronic changes in dry heat transfer (q_Tot_) at the eye region of black-capped chickadees (n = 19) following stress exposure across varying ambient temperatures. Individual lines represents the predicted correlation between ambient temperature (here, mean-centered) and q_Tot_ of individual black-capped chickadees during stress exposure or control treatments. Grey rectangle represents the thermoneutral zone (TNZ) for black-capped chickadees (estimated from Grossman and West, 1977). Bold black lines (solid and dashed) and accompanying delta (*δ*) symbols indicate the spread of correlations between ambient temperature and q_Tot_ across individuals, in control and stress exposure treatments respectively. Correlations are estimated from a Bayesian generalised additive mixed effects model (GAMM) and marginalised across all environmental and experimental parameters. q_Tot_ values were estimated by infra-red thermography (n = 5832 images) across 60 days.

As predicted, acute stress-induced changes in T_s_ and q_Tot_ (or “acute reaction norms”) were significantly repeatable among chickadees. Namely, repeatability values calculated from our true models exceeded those calculated from our null models (i.e. with individual identities scrambled; non-linear hypothesis test: K_Ts_ > 100; K_qTot_ = 47.00; Figure 5a and Figure S5a), suggesting that repeatability of acute thermal responses to stress exposure not only exceeded zero, but also could not be explained by biases in our experimental methodology. Nevertheless, the degree to which these acute thermal responses were repeatable among chickadees was low (surface temperature [T_s_]: R_stress exposure_ = 0.14 [0.03, 0.32]; heat transfer [q_Tot_]: R_stress exposure_ = 0.11 [0.02, 0.27]; Table 1), suggesting that while some variation in acute thermal responses is probably attributable to consistent differences in stress-responsive phenotypes among individuals, the majority of such variation is perhaps better explained by other sources of variation (e.g. environmental or measurement). Similar to acute changes in T_s_ and q_Tot_, chronic changes in T_s_ and q_Tot_ following stress exposure (or “chronic reaction norms”) were significantly repeatable among chickadees. Again, repeatability values estimated from our true models exceeded those estimated from our null models, suggesting that repeatability of chronic changes in T_s_ and q_Tot_ observed in our study were unlikely to be explained by biases in our experimental method (non-linear hypothesis tests comparing true and null models; K > 100 for both T_s_ and q_Tot_; Figure 5b and Figure S5b). Here, however, repeatability of chronic reaction norms among chickadees was high (R_Ts_ = 0.61 [0.35, 0.81]; R_qTot_ = 0.67 [0.44, 0.84]; Table 2), indicating that long-term stress-induced changes in T_s_ and q_Tot_ consistently varied among individuals.

**FIGURE 5.**
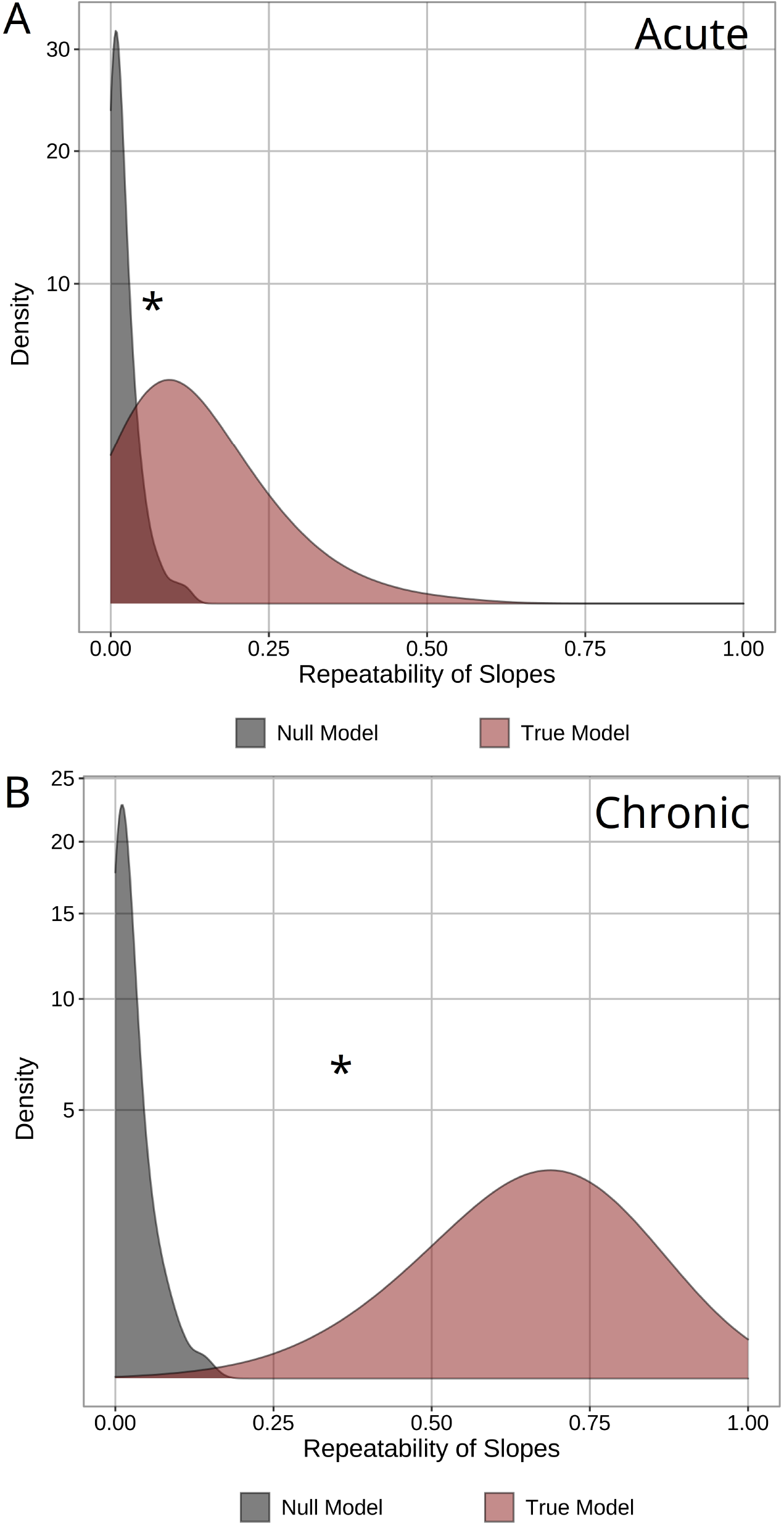
Repeatability of acute and chronic changes in dry heat transfer (q_Tot_) at the eye region during stress exposure in black-capped chickadees (n = 19). Panels **A** and **B** represent distribution of repeatability values for acute and chronic responses to stress exposure, respectively. True model distributions (red) represent those of drawn from models where identity of individuals was correctly identified. In contrast, null model distributions (grey) represent those drawn from models where identity of individuals was randomly scrambled. A positive difference between true and null distributions (indicated by an asterisk, “*”) implies that repeatability values from true models cannot be explained by biases in experimental methods (captured in null models) and are considered significant. Distributions are estimated from posteriors of Bayesian generalised additive mixed effects models (GAMM). Thermal responses to stress exposure represent those observed at the eye region of chickadees, using infra-red thermography across 60 days of observation.

### 3.2 Evidence for stabilising or directional selection on stress-induced changes in body surface temperature and peripheral heat loss

Across acute time-periods (i.e. ≤ 1 hour), T_s_ of control individuals was significantly more variable and less consistent than that of stress-exposed individuals, after controlling for circadian rhythms and environmental effects (e.g. ambient temperature, solar radiation; *σ*_Control_ = 1.21 [1.19, 1.24], *σ*_Stress_ = 1.18 [1.14, 1.22]; R_control_ = 0.07 [0.01, 0.18], R_stress exposure_ = 0.14 [0.03, 0.32]; Table 4.1). As predicted, these difference in variance and repeatability between treatments were strongly and moderately supported by non-linear hypothesis tests respectively (K_variance_ = 72.47; K_repeatability_ = 6.66; Figure S6). Similarly, q_Tot_ at the eye region of chickadees was both slightly less variable and more repeatable during stress exposure treatments than control treatments (*σ*_Control_ = 2.09 [2.06, 2.13], *σ*_Stress_ = 2.06 [1.99, 2.12]; R_control_ = 0.06 [0.01, 0.16], R_stress exposure_ = 0.11 [0.02, 0.27]; Table 1). These differences in unexplained variability and repeatability, however, were only moderately and weakly supported by non-linear hypothesis tests respectively (K_variance_ = 10.65; K_repeatability_ = 5.24; Figure S7).

Similar to acute time periods, T_s_ of chickadees was more variable and less repeatable in control treatments than in stress exposure treatments across chronic time periods (i.e. ≤ 30 days), after controlling for circadian and environmental effects (*σ*_Control_ = 1.20 [1.18, 1.23], *σ*_Stress_ = 1.17 [1.13, 1.21]; R_control_ = 0.34 [0.17, 0.56], R_stress exposure_ = 0.61 [0.35, 0.81]; Table 2). Again, as predicted, these differences in variance and repeatability were strongly and moderately supported by respective non-linear hypothesis tests (K_variance_ = 48.32; K_repeatability_ = 15.51; Figure S8). Variability and repeatability of q_Tot_ across chronic time periods followed similar patterns, with variability again being lower and repeatability again being higher in stress-exposed chickadees, when compared with rested (i.e. control) chickadees (*σ*_Control_ = 2.07 [2.04, 2.11], *σ*_Stress_ = 2.04 [1.98, 2.10]; R_control_ = 0.41 [0.22, 0.63], R_stress exposure_ = 0.67 [0.44, 0.84]; Table 2). These differences were moderately and strongly supported by non-linear hypothesis tests respectively (K_variance_ = 9.62; K_repeatability_ = 16.73; Figure S9), as predicted.

### 3.3 Stress-induced thermal responses do not differ between urban and rural individuals

The magnitude of acute changes in T_s_ or q_Tot_ (or “acute reaction norms”) following stress exposure did not differ between chickadees captured from urban or rural ecotypes (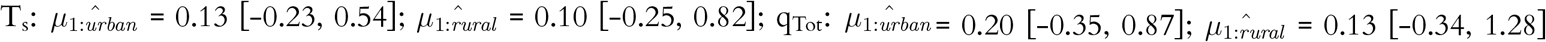; n = 9 urban, n = 10 rural; Figure 6; Figure S10). Indeed, ANOVAs including capture ecotype as a population-level predictor were less likely to explain the magnitude of T_s_ or q_Tot_ responses among individuals than ANOVAs without (T_s_: K = 0.24; q_Tot_: K = 0.25). Similar results were detected at the chronic level, with the magnitude of chronic stress-induced changes in T_s_ and q_Tot_ (or, “chronic reaction norms”) remaining similar between urban- and rural-origin chickadees (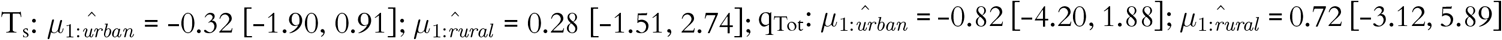; n = 9 urban, n = 10 rural; Figure 6). Again, ANOVAs including capture ecotype as a predictor were less likely to explain the magnitudes of chronic changes in T_s_ and q_Tot_ than ANOVAs without (T_s_: K = 0.39; q_Tot_: K = 0.49).

**FIGURE 6.**
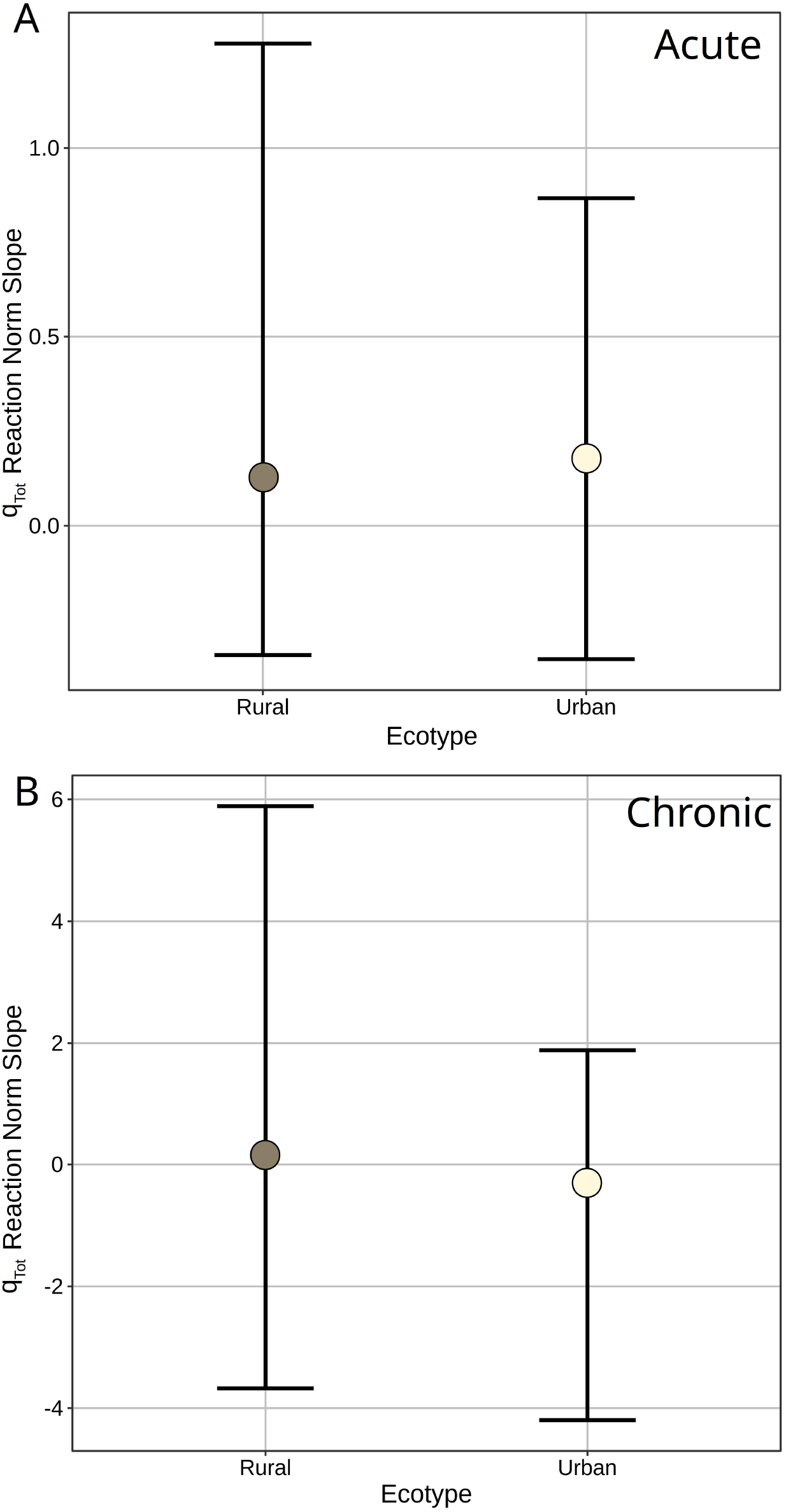
Average effect of stress exposure on dry heat transfer (q_Tot_; reaction norm slopes) at the eye region of black-capped chickadees (n = 19) captured from urban and rural ecotypes (n = 9 urban, n = 10 rural). **A** | Average slopes of acute reaction norm across individuals captured at each ecotype. Reaction norm slopes represent the slopes of the linear interaction between treatment type and time post stress exposure (s) per individual. **B** | Average slopes of chronic reaction norms across individuals captured from each ecotype. Here, reaction norm slopes represent those of linear interactions between treatment type and ambient temperature (°C) per individual. Error bars represent 95% credible intervals around mean estimates. All reaction norm slopes were derived from Bayesian generalised additive mixed effects models (GAMMs).

## 4 DISCUSSION

### 4.1 Acute and chronic thermal responses to stress exposure are repeatable

Our results show that flexible changes in surface temperature (T_s_) and rate of heat transfer (q_Tot_) following stress exposures are repeatable in chickadees, whether observed across acute or protracted (i.e. chronic) time periods. Such repeatability fulfills a critical first prediction of the hypothesis that stress-induced flexibility of T_s_ and q_Tot_ may experience evolutionary responses to selection. Notably, however, the extent to which flexibility of T_s_ and q_Tot_ was repeatable appeared to depend upon the time period of observation (Figure 5 and Figure S5). Across acute time periods, the shape and magnitude of stress-induced T_s_ and q_Tot_ responses were appreciably similar among individuals (Figures 3b and 5a; Figure S5a). Across chronic time-periods, however, a considerably wider range of stress-responsive phenotypes among individuals emerged (Figures 4 and 5b; Figure S5b). To our knowledge, our study is the first to report repeatability of stress-induced flexibility of T_s_ and q_Tot_ in any vertebrate.

The high degree with which chronic responses to stress exposure varied among our study individuals highlights that, despite a clear average trend among individuals (Figure 4; Table 1), reductions in average T_s_ and q_Tot_ in the cold and increases in average T_s_ and q_Tot_ in the warmth are clearly not generalisable responses to repeated stress perception in birds. Among some individuals, for example, repeated stress exposure appeared to elicit the reverse response, with mean T_s_ and q_Tot_ rising in the cold and decreasing in the heat (Figure 4). If the emergence of such chronic stress-induced responses are largely fixed within individuals, as our study suggests, theorised energetic benefits ascribed to this response (e.g. Robertson *et al*. 2020b; Jerem *et al*. 2018; Herborn *et al*. 2018) may only be accrued by some and not all individuals. Given that survivorship has been linked to efficiency of energy use in extreme and challenging environments (Parsons, 2005), such discrepancies in theorised energetic savings could provide opportunities for selection to act upon chronic thermal responses to stress exposure in our study species.

Any evolutionary responses to selection on flexibility of T_s_ and q_Tot_ in response to chronic stress exposures requires that this trait is underpinned by heritable genetic architecture. In this study, we chose to monitor changes in T_s_ and peripheral q_Tot_ in response to stress exposure alone. Therefore, whether chronic responses observed here emerge as a consequence of stress-induced changes in core body temperature, peripheral temperature (e.g. by changes in vascular flow; Oka *et al*. 2001), or both remains unknown. Regardless of their anatomical origin, the possibility of individual differences in chronic responses arising from differences in genetic architecture is well supported. At the level of core tissues, for example, both heterothermy and facultative hypothermia appear phylogenetically constrained (Boyles *et al*., 2013; Gerson *et al*., 2019), and recent studies in poultry have provided strong evidence for the direct influence of genetic polymorphisms and differential gene transcription on heat dissipation capacity and the magnitude of core body temperature increases in suprathermoneutral ambient temperatures (Srikanth *et al*., 2019; Zhuang *et al*., 2019). Similarly, at the level of the periphery, studies in humans have elucidated several genetic polymorphisms that appear to dictate the duration and magnitude of peripheral vascular responses to cold and psychological stress (e.g. Rao *et al*. 2008; Chen *et al*. 2010; Kelsey *et al*. 2010, 2012; Huang *et al*. 2012) that could have meaningful consequences on environmental heat transfer; many such polymorphisms correspond to genes with conserved functions among tetrapods (Vincent *et al*. 1998; Yamamoto & Vernier 2011; Céspedes *et al*. 2017; Dopamine *β*-hydroxylase in sauropsids: Lovell *et al*. 2015). Consequently, variation in stress-induced changes in T_s_ and q_Tot_ among our chickadees may well be heritable, regardless of whether such responses are driven by changes in thermogenesis at the core, or by changes in peripheral vascular flow and consequential changes in environmental heat transfer.

Still, we cannot refute the possibility that our observed chronic responses to stress exposure are broadly labile within individuals and dictated by energetic or resource constraints that were not measured here. For example, Robertson et al (2020b) recently argued that stress-induced changes in T_s_ and q_Tot_ may be understood as trade-offs that are predominantly manifested under negative energetic balance (see Oka 2018 and suggestions by Lewden *et al*. 2017; Winder *et al*. 2020). It is possible that our experimental conditions may have contributed to fixed and non-random resource allocation among individuals (e.g. via dominance interactions; Ratcliffe *et al*. 2007) that dictated how stress-induced thermal responses at the eye region emerged. In such a case, any evolutionary responses to selection on stress-induced thermal flexibility may better reflect patterns of resource monitoring and allocation during a challenge, rather than fixed reflexes within individuals. Although our results suggest that the repeatabilities of both acute and chronic stress-induced thermal responses are unlikely to be explained by variations in resource access (Supporting Information; Figures S11-S12), futher experiments seeking to tease apart the influence of resource availability and fixed individual variation on chronic thermal responses to stress exposure are therefore warranted.

To our surprise, the degree to which individuals acutely shifted their T_s_ and q_Tot_ in response to stress exposures displayed considerable overlap (Figures 3a and 3b). Such overlap among individuals, coupled with the significant predictive effects of other environmental parameters (e.g. ambient temperature and time of day; Table 1) implies that, unlike chronic thermal responses, the manifestation of acute thermal responses to stress exposure is perhaps better explained by the combination of common trait expression and environmental effects than variation in intrinsic factors among individuals. In domestic rats (*Rattus norvegicus domestica*), ambient temperature has been shown to strongly influence the magnitude of acute changes in core body temperature, with responses typically being largest at low ambient temperature and smallest at high ambient temperatures (Briese 1992; reviewed in Oka 2018). Similarly, in Svalbard rock ptarmigans (*Lagopus muta hyperborea*), the magnitude of stress-induced changes in skin temperature are reportedly larger at low ambient temperature than at comparatively higher ambient temperatures (Nord & Folkow, 2019). As such, the emergence of acute, stress-induced changes in T_s_ and q_Tot_ in our sample population may have been largely dictated by modulatory effects of ambient temperature alone, with little remaining variation explained by phenotypic differences among individuals. In any case, the relatively low repeatability of acute stress-induced thermal responses (observed here) highlights that the potential for this response to respond to selection in black-capped chickadees is probably low.

### 4.2 Variation in eye region temperature and heat loss is reduced during stress exposure

Interestingly, unexplained variation in both T_s_ and q_Tot_ was higher during control treatments than during stress exposure treatments (Tables 1-2). Additionally, both T_s_ and q_Tot_ were more repeatable during stress exposure treatments than control treatments (Figures S6-S9) regardless of the time period of observation (i.e. ≤ 1 hour, or ≤ 30 days). Together, these trends indicate that either: (1) T_s_ and q_Tot_ are more tightly regulated during stress exposures than during resting conditions, or (2) T_s_ regulation is relaxed during stress exposures, thereby allowing T_s_ to conform to ambient temperatures (as observed in other avian species; reviewed in Angilletta *et al*. 2019). Regardless of the mechanism, the relative consistency with which T_s_ and q_Tot_ emerge during stress exposures suggests that their manifestation has, perhaps, experienced stronger stabilising or directional selection than that during rested (i.e. control) conditions (our second prediction; e.g. Gibson & Bradley 1974; Lande & Arnold 1983; Van Homrigh *et al*. 2007; but see Kotiaho *et al*. 2001). Such findings lend credence to a critical role of heat-transfer regulation during stress exposure, that, to our knowledge, has received little to no research attention.

When contextualised with variability of other stress-physiological processes, reduced variability of T_s_ and q_Tot_ during stress perception is perhaps not unusual. Variability in heart rate is widely known to fall during stress exposure in many vertebrate species (e.g. Visser *et al*. 2002; Von Borell *et al*. 2007; Cyr *et al*. 2009). Similarly, within-individual variation in stress-induced glucocortoid production has been reported to be lower than that of baseline production in both avian and amphibian species (e.g. Cockrem & Silverin 2002; Rensel & Schoech 2011; Narayan *et al*. 2012; Grace & Anderson 2014; but see Narayan *et al*. 2013; Baugh *et al*. 2014; Lendvai *et al*. 2015). Such trends indicate that the collective traits enabling individuals to conform or cope with environmental challenges (together, the “stress phenotype”) have experienced strong stabilising or directional selection (Ellis *et al*., 2006). Modulation of T_s_ and q_Tot_ during stress exposure (whether by a reduction or increase) may, therefore, simply represent a little-discussed constituent of the vertebrate stress phenotype that contributes to successful coping. Although the ultimate value of stress-induced T_s_ and q_Tot_ modulation is unclear, the bivalent nature, ambient-temperature dependence, and direct implications on energetic savings in our study (albeit small; Figure 4) triangulate on a relaxation of expenditure toward thermoregulation (the Thermoprotective Hypothesis; Robertson *et al*. 2020a). On the other hand, rapid increases in T_s_ and q_Tot_ at low ambient temperatures, and rapid declines in T_s_ and q_Tot_ following stress exposure (as observed here; Figures 3a and 3b) may suggest that at the acute level, changes in T_s_ occur to promote enzymatic, neuronal, or muscular function during the stress responses (i.e. owing to Q10 effects: e.g. Carr & Lima 2013), rather than to reduce thermoregulatory expenses.

### 4.3 Urban and rural individual do not differ in stress-induced thermal responses

In sharp contrast to our predictions, the degree to which T_s_ and q_Tot_ flexibly responded to acute or chronic stress exposure did not differ between chickadees captured from urban and rural environments (Figure 6 and Figure S10). According to our results, individuals from urban environments appear no more able to flexibly shift their T_s_ and thermoregulatory expenditure during stress exposure than those from rural environments. We propose four possible explanations for these findings. First, insufficient generations spent within a given ecotype may have limited opportunities for evolutionary responses to selection on stress-induced thermal responses to occur in our study species. The combination of low juvenile dispersal, high site fidelity among adults (Weise & Meyer, 1979), and relatively short generation time in our study species, however, suggests that this is unlikely (reviewed in McDonnell & Hahs 2015). Furthermore, genetic differentiation between individuals captured in urban and rural environments has recently been reported for a closely related Parid species (the great tit, *Parus major*; Perrier *et al*. 2018), supporting the possibility of responses to selection imposed by urban environments. A second, and arguably more likely explanation for our findings is that costs of urban living in chickadees are no higher than those of rural living, despite a theoretically increased frequency in stress exposure events. Although direct comparative field studies are lacking (Sepp *et al*., 2018), trends in basal metabolic rate of another temperature bird species (the house finch, *Haemorhous mexicanus*) do suggest that energetic expenditure may not differ between individuals captured from urban and rural environments (at least, at rest: Hutton *et al*. 2018). In chickadees, urban environments may afford opportunities to access novel and abundant food sources (Robb *et al*., 2008; Prasher *et al*., 2019) that could offset energetic costs associated with frequent activation of emergency pathways (but see Demeyrier *et al*. 2017). Strategies to relax expenditure towards other biological process (e.g. thermoregulation), therefore, may be no more likely to emerge in urban population than rural populations. Third, the degree of urbanisation in our selected urban and rural locations may not have differed sufficiently to impose differential patterns of selection (but see Appendix). Given our low sample size, assessing linear correlations between the degree of urbanisation at capture locations and the magnitude of stress-induced thermal responses was unfortunately not possible. Future studies assessing these response among individuals from more urbanised locations may be warranted. Lastly, neither acute nor chronic changes in T_s_ and q_Tot_ that accompany stress exposures may be heritable in chickadees. Previous studies, both within and across species, have suggested that changes in core body temperature and peripheral vascular flow during a challenge are underpinned by heritable genetic architecture (discussed above). Nevertheless, it is indeed possible that thermal responses to stress exposure at either the acute or chronic level are merely contingent upon environmental context (e.g. resource availability) and the maximum degree to which T_s_ and q_Tot_ can flexibly respond to stress exposure is fixed among individuals. Further studies questioning the heritability of stress-induced thermal responses in this species are, therefore, critical to understanding whether this response may provide opportunities to adapt to a warming and urbanising world.

### 4.4 Summary

Recent empirical studies have argued that endotherms may balance costs associated with responding to perceived stressors by flexibly decreasing their T_s_ and q_Tot_ in the cold, and flexibly increasing their T_s_ and q_Tot_ in warmth. By doing so, energy may be allocated away from costly thermogenesis or evaporative cooling, and toward the immediate demands of coping with the challenge at hand. In chickadees, we tested whether such stress-induced flexibilities of T_s_ and q_Tot_ are repeatable among individuals and thus offer opportunities for endotherms to cope with costs that typify urbanised environments, across generations. As predicted, we show that both acute and chronic changes in T_s_ and q_Tot_ during stress exposure are repeatable, however, only those at the chronic level displayed meaningfully high repeatability estimates (T_s_: R_chronic_ = 0.61; q_Tot_: R_chronic_ = 0.67). Furthermore, we show that both T_s_ and q_Tot_ are less variable within individuals, and more variable among individuals during experimental stress exposure than during control treatment, suggesting that regulation of T_s_ and q_Tot_ during the stress response has probably experienced stabilising or directional selection. Both trends, to our knowledge, are yet to be reported in any vertebrate. To our surprise, neither acute, nor chronic flexibility of T_s_ and q_Tot_ in response to stress exposure differed between urban- and rural-origin chickadees. Together, our results suggest that while flexibility of T_s_ and q_Tot_ meet a critical first criterion for responsiveness to selection and may enhance energetic efficiency of some but not all individuals, those residing in urban environments are no more likely to acquire benefits associated with this flexibility than those in rural environments.

## Acknowledgments

First, we thank both Lianne Ralph and the collective staff at the Ruthven Park National Historic Site for their assistance in experimental execution, without which this study would not have been possible. Furthermore, we thank Kimberley Tasker, Simon Tapper and Tyler Maksymiw for their assistance in aviary construction, Simon Tapper for his statistical insight, and Elise Cote for her artistic contributions to figures 1 and 2. This research was supported by an NSERC Discovery grant (RGPIN-04158-2014) to GB and an NSERC CREATE grant (RGPIN-481954-2016) to GB and GM.

## References

Andreasson, F., Nord, A. & Nilsson, J. (2020) Body temperature responses of great tits parus major to handling in the cold. Ibis 162(3), 836–844.

Angel, S., Parent, J., Civco, D., Blei, A. & Potere, D. (2011) The dimensions of global urban expansion: Estimates and projections for all countries, 2000-2050. Progress in Planning 75(2), 53–107.

Angilletta, M., Youngblood, J., Neel, L. & VandenBrooks, J. (2019) The neuroscience of adaptive thermoregulation. Neuroscience Letters 692(1), 127–136.

Araya-Ajoy, Y., Mathot, K. & Dingemanse, N. (2015) An approach to estimate short-term, longterm and reaction norm repeatability. Methods in Ecology and Evoloution 6(12), 1462–1473.

Argüeso, D., Evans, J., Pitman, A. & Di Luca, A. (2015) Effects of city expansion on heat stress under climate change conditions. PloS One 10(2), e0117066.

Arnfield, A. (2003) Two decades of urban climate research: a review of turbulence, exchanges of energy and water, and the urban heat island. International Journal of Climatology 23(1), 1–26.

Barton, A., Irwin, A., Finkel, Z. & Stock, C. (2016) Anthropogenic climate change drives shift and shuffle in north atlantic phytoplankton communities. Proceedings of the National Academy of Sciences 113(11), 2964–2969.

Baugh, A., van Oers, K., Dingemanse, N. & Hau, M. (2014) Baseline and stress-induced glucocorticoid concentrations are not repeatable but covary within individual great tits (parus major). General and Comparative Endocrinology 208(1), 154–163.

Best, R. & Fowler, R. (1981) Infrared emissivity and radiant surface temperatures of Canada and snow geese. Journal of Wildlife Management 45(4), 1026–1029.

Birnie-Gauvin, K., Peiman, K., Gallagher, A., De Bruijn, R. & Cooke, S. (2016) Sublethal consequences of urban life for wild vertebrates. Environmental Reviews 24(1), 416–425.

Blair, D., Glover, W., Greenfield, A. & Roddie, I. (1959) Excitation of cholinergic vasodilator nerves to human skeletal muscles during emotional stress. Journal of Physiology 148(3), 633.

Boake, C. (1989) Repeatability: its role in evolutionary studies of mating behavior. Evolutionary Ecology 3(2), 173–182.

Bonier, F. (2012) Hormones in the city: endocrine ecology of urban birds. Hormones and Behavior 61(5), 763–772.

Boratynski, J., Iwińska, K. & Bogdanowicz, W. (2019) An intra-population heterothermy continuum: notable repeatability of body temperature variation in food-deprived yellow-necked mice. Journal of Experimental Biology 222(6), jeb197152.

Boyles, J., Thompson, A., McKechnie, A., Malan, E., Humphries, M. & Careau, V. (2013) A global heterothermic continuum in mammals. Global Ecology and Biogeography 22(1), 1029–1039.

Brans, K., Jansen, M., Vanoverbeke, J., Tüzün, N., Stoks, R. & De Meester, L. (2017) The heat is on: genetic adaptation to urbanization mediated by thermal tolerance and body size. Global Change Biology 23(12), 5218–5227.

Breuner, C. & Berk, S. (2019) Using the van Noordwijk and de Jong resource frame-work to evaluate glucocorticoid-fitness hypotheses. Integrative and Comparative Biology 59(2), 243–250.

Briese, E. (1992) Cold increases and warmth diminishes stress-induced rise of colonic temperature in rats. Physiology and Behavior 51(4), 881–883.

Bürkner, P. (2017) brms: An r package for bayesian multilevel models using stan. Journal of Statistical Software 80(1), 1–28.

Careau, V., Réale, D., Garant, D., Speakman, J. & Humphries, M. (2012) Stress-induced rise in body temperature is repeatable in free-ranging eastern chipmunks (tamias striatus). Journal of Comparative Physiology B 182(3), 403–414.

Carr, J. & Lima, S. (2013) Nocturnal hypothermia impairs flight ability in birds: a cost of being cool. Proceedings of the Royal Society of London B: Biological Sciences 280(1772), 20131846.

Céspedes, H., Zavala, K., Vandewege, M. & Opazo, J. (2017) Evolution of the alpha2-adrenoreceptors in vertebrates: ADRA2D is absent in mammals and crocodiles. General and Comparative Endocrinology 250(1), 85–94.

Chen, Y., Wen, G., Rao, F., Zhang, K., Wang, L., Rodriguez-Flores, J., Sanchez, A., Mahata, M., Taupenot, L., Sun, P. & Mahata, S. (2010) Human dopamine beta-hydroxylase (DBH) regulatory polymorphism that influences enzymatic activity, autonomic function, and blood pressure. Journal of Hypertension 28(1), 76.

Cockrem, J. & Silverin, B. (2002) Variation within and between birds in corticosterone responses of great tits (parus major). General and Comparative Endocrinology 125(2), 197–206.

Cooper, S. & Gessaman, J. (2005) Nocturnal hypothermia in seasonally acclimatized mountain chickadees and juniper titmice. Condor 107(1), 151–155.

Cyr, N., Dickens, M. & Romero, L. (2009) Heart rate and heart-rate variability responses to acute and chronic stress in a wild-caught passerine bird. Physiology and Biochemical Zoology 82(4), 332–344.

de Aguiar Bittencourt, M., Melleu, F. & Marino-Neto, J. (2015) Stress-induced core temperature changes in pigeons (columba livia). Physiology and Behavior 139(1), 449–458.

Demeyrier, V., Charmantier, A., Lambrechts, M. & Grégoire, A. (2017) Disentangling drivers of reproductive performance in urban great tits: a food supplementation experiment. Journal of Experimental Biology 220(22), 4195–4203.

Depke, M., Fusch, G., Domanska, G., Geffers, R., Völker, U., Schuett, C. & Kiank, C. (2008) Hypermetabolic syndrome as a consequence of repeated psychological stress in mice. Endocrinology 149(6), 2714–2723.

Dochtermann, N., Schwab, T. & Sih, A. (2015) The contribution of additive genetic variation to personality variation: heritability of personality. Proceedings of the Royal Society of London B: Biological Sciences 282(1), 20142201.

Dohm, M. (2002) Repeatability estimates do not always set an upper limit to heritability. Functional Ecology 16(1), 273–280.

du Plessis, K., Martin, R., Hockey, P., Cunningham, S. & Ridley, A. (2012) The costs of keeping cool in a warming world: implications of high temperatures for foraging, thermoregulation and body condition of an arid zone bird. Global Change Biology 18(10), 3063–3070.

Ellis, B., Jackson, J. & Boyce, W. (2006) The stress response systems: universality and adaptive individual di?erences. Developmental Review 26(2), 175–212.

Fokidis, H., Orchinik, M. & Deviche, P. (2009) Corticosterone and corticosteroid binding globulin in birds: relation to urbanization in a desert city. General and Comparative Endocrinology 160(3), 259–270.

Freeman, L., Corbett, D., Fitzgerald, A., Lemley, D., Quigg, A. & Steppe, C. (2019) Impacts of urbanization and development on estuarine ecosystems and water quality. Estuaries and Coasts 42(7), 1821–1838.

French, S., Fokidis, H. & Moore, M. (2008) Variation in stress and innate immunity in the tree lizard (urosaurus ornatus) across an urban-rural gradient. Journal of Comparative Physiology B 178(8), 997–1005.

Gerson, A., McKechnie, A., Smit, B., Whitfield, M., Smith, E., Talbot, W., McWhorter, T. & Wolf, B. (2019) The functional significance of facultative hyperthermia varies with body size and phylogeny in birds. Functional Ecology 33(4), 597–607.

Gibson, J. & Bradley, B. (1974) Stabilising selection in constant and fluctuating environments. Heredity 33(3), 293–302.

Grace, J. & Anderson, D. (2014) Corticosterone stress response shows long-term repeatability and links to personality in free-living Nazca boobies. General and Comparative Endocrinology 208(1), 39–48.

Grimm, N., Faeth, S., Golubiewski, N., Redman, C., Wu, J., Bai, X. & Briggs, J. (2008) Global change and the ecology of cities. Science 319(5864), 756–760.

Grossman, A. & West, G. (1977) Metabolic rate and temperature regulation of winter acclimatized black-capped chickadees Parus atricapillus of interior alaska. Ornis Scandinavica 8(2), 127–138.

Harris, A. (2016) Fitsio. R package, version 2.1-0.

Herborn, K., Jerem, P., Nager, R., McKeegan, D. & McCafferty, D. (2018) Surface temperature elevated by chronic and intermittent stress. Physiology and Behavior 191(1), 47–55.

Hernández-Brito, D., Carrete, M., Popa-Lisseanu, A., Ibáñez, C. & Tella, J. (2014) Crowding in the city: losing and winning competitors of an invasive bird. PloS One 9(6), e100593.

Hill, R., Beaver, D. & Veghte, J. (1980) Body surface temperatures and thermoregulation in the black-capped chickadee (parus atricapillus). Physiological Zoology 53(3), 305–321.

Huang, J., Chen, S., Lu, X., Zhao, Q., Rao, D., Jaquish, C., Hixson, J., Chen, J., Wang, L., Cao, J. & Li, J. (2012) Polymorphisms of ACE2 are associated with blood pressure response to cold pressor test: the GenSalt study. American Journal of Hypertension 25(8), 937–942.

Hutton, P., Wright, C., DeNardo, D. & McGraw, K. (2018) No e?ect of human presence at night on disease, body mass, or metabolism in rural and urban house finches (haemorhous mexicanus). Integrative Comparative Biology 58(5), 977–985.

Ikkatai, Y. & Watanabe, S. (2015) Eye surface temperature detects stress response in budgerigars (Melopsittacus undulatus). NeuroReport 26(11), 642–646.

Jerem, P., Herborn, K., McCafferty, D., McKeegan, D. & Nager, R. (2015) Thermal imaging to study stress non-invasively in unrestrained birds. Jouranl of Visualized Experiments (JoVE) pp. 1–10.

Jerem, P., Jenni-Eiermann, S., Herborn, K., McKeegan, D., McCafferty, D. & Nager, R. (2018) Eye region surface temperature reflects both energy reserves and circulating glucocorticoids in a wild bird. Scientific Reports 8(1), 1907.

Jerem, P., Jenni-Eiermann, S., McKeegan, D., McCafferty, D. & Nager, R. (2019) Eye region surface temperature dynamics during acute stress relate to baseline glucocorticoids independently of environmental conditions. Physiology and Behavior 210(1), 112627.

Jimeno, B., Hau, M. & Verhulst, S. (2017) Strong association between corticosterone levels and temperature-dependent metabolic rate in individual zebra finches. Journal of Experimental Biology 220(23), 4426–4431.

Johnson, J., Trubl, P. & Miles, L. (2012) Black widows in an urban desert: city-living compromises spider fecundity and egg investment despite urban prey abundance. The American Midland Naturalist 168(2), 333–340.

Kelsey, R., Alpert, B., Dahmer, M., Krushkal, J. & Quasney, M. (2010) Betaadrenergic receptor gene polymorphisms and cardiovascular reactivity to stress in black adolescents and young adults. Psychophysiology 47(5), 863–873.

Kelsey, R., Alpert, B., Dahmer, M., Krushkal, J. & Quasney, M. (2012) Alphaadrenergic receptor gene polymorphisms and cardiovascular reactivity to stress in black adolescents and young adults. Psychophysiology 49(3), 401–412.

Knauer, J., El-Madany, T., Zaehle, S. & Migliavacca, M. (2018) Bigleaf - an R package for the calculation of physical and physiological ecosystem properties from eddy covariance data. PLoS One 13(8), e0201114.

Kotiaho, J., Simmons, L. & Tomkins, J. (2001) Towards a resolution of the lek paradox. Nature 410(6829), 684–686.

Lande, R. & Arnold, S. (1983) The measurement of selection on correlated characters. Evolution 37(6), 1210–1226.

Lendvai, Á., Giraudeau, M., Bókony, V., Angelier, F. & Chastel, O. (2015) Within-individual plasticity explains age-related decrease in stress response in a short-lived bird. Biology Letters 11(7), 20150272.

Lerch, M. (2017) International migration and city growth. Population Division Technical Paper 2017/10, United Nations, New York, USA.

Lewden, A., Nord, A., Petit, M. & Vézina, F. (2017) Body temperature responses to handling stress in wintering black-capped chickadees (poecile atricapilus). Physiology and Behavior 179(1), 49–54.

Lovell, P., Kasimi, B., Carleton, J., Velho, T. & Mello, C. (2015) Living without DAT: Loss and compensation of the dopamine transporter gene in sauropsids (birds and reptiles). Scientific Reports 5(1), 1–12.

Lowry, H., Lill, A. & Wong, B. (2013) Behavioural responses of wildlife to urban environments. Biological Reviews 88(3), 537–549.

Mainwaring, M., Barber, I., Deeming, D., Pike, D., Roznik, E. & Hartley, I. (2017) Climate change and nesting behaviour in vertebrates: a review of the ecological threats and potential for adaptive responses. Biological Reviews 92(4), 1991–2002.

McCafferty, D., Gilbert, C., Paterson, W., Pomeroy, P., Thompson, D., Currie, J. & Ancel, A. (2011) Estimating metabolic heat-loss in birds and mammals by combining infrared thermography with biophysical modelling. Comparative Biochemistry and Physiology A: Molecular & Integrative Physiology 158(3), 337–345.

McDonnell, M. & Hahs, A. (2015) Adaptation and adaptedness of organisms to urban environments. Annual Review of Ecology, Evolution, and Systematics 46(1), 261–280.

Minkina, W. & Dudzik, S. (2009) Infrared thermography errors and uncertainties. Wiley Press, Chichester, UK.

Morey, R., Rouder, J., Jamil, J. & Morey, M. (2019) Bayesfactor. R package, version 0.9.12.4.2.

Müller, M., Vyssotski, A., Yamamoto, M. & Yoda, K. (2018) Individual di?erences in heart rate reveal a broad range of autonomic phenotypes in a free-living seabird population. Journal of Experimental Biology 221(19), jeb182758.

Narayan, E., Cockrem, J. & Hero, J. (2013) Are baseline and short-term corticosterone stress responses in free-living amphibians repeatable? Comparative Biochemistry and Physiology A: Molecular & Integrative Physiology 164(1), 21–28.

Narayan, E., Molinia, F., Cockrem, J. & Hero, J. (2012) Individual variation and repeatability in urinary corticosterone metabolite responses to capture in the cane toad (rhinella marina). General and Comparative Endocrinology 175(2), 284–289.

Nespolo, R. & Franco, M. (2007) Whole-animal metabolic rate is a repeatable trait: a meta-analysis. Journal of Experimental Biology 210(11), 2000–2005.

Newsome, S., Garbe, H., Wilson, E. & Gehrt, S. (2015) Individual variation in anthropogenic resource use in an urban carnivore. Oecologia 178(1), 115–128.

Nord, A. & Folkow, L. (2019) Ambient temperature effects on stress-induced hyper-thermiain Svalbard ptarmigan. Biology Open 8(6), bio043497.

Nord, A. & Nilsson, J. (2019) Heat dissipation rate constrains reproductive investment in a wild bird. Functional Ecology 33(2), 250–259.

Oka, T. (2018) Stress-induced hyperthermia and hypothermia. Handbook of Clinical Neurology (ed. A. Romanovsy), vol. 157, pp. 599–621, Elsevier, Amsterdam, Netherlands.

Oka, T., Oka, K. & Hori, T. (2001) Mechanisms and mediators of psychological stress-induced rise in core temperature. Psychosomatic Medicine 63(1), 476–486.

Ouyang, J., Hau, M. & Bonier, F. (2011) Within seasons and among years: when are corticosterone levels repeatable? Hormones and Behavior 60(5), 559–564.

Ouyang, J., Isaksson, C., Schmidt, C., Hutton, P., Bonier, F. & Dominoni, D. (2018) A new framework for urban ecology: an integration of proximate and ultimate responses to anthropogenic change. Integrative and Comparative Biology 58(5), 915–928.

Pacifici, M., Visconti, P., Butchart, S., Watson, J., Cassola, F. & Rondinini, C. (2017) Species’ traits influenced their response to recent climate change. Nature Climate Change 7(3), 205–208.

Parsons, P. (2005) Environments and evolution: interactions between stress, resource inadequacy and energetic e?ciency. Biological Reviews 80(4), 589–610.

Partecke, J., Schwabl, I. & Gwinner, E. (2006) Stress and the city: urbanization and its e?ects on the stress physiology in European blackbirds. Ecology 87(8), 1945–1952.

Pautasso, M. (2012) Observed impacts of climate change on terrestrial birds in Europe: an overview. Italian Journal of Zoology 79(2), 296–314.

Pendlebury, C., MacLeod, M. & Bryant, D. (2004) Variation in temperature increases the cost of living in birds. Journal of Experimental Biology 207(12), 2065–2070.

Perrier, C., Lozano del Campo, A., Szulkin, M., Demeyrier, V., Gregoire, A. & Charmantier, A. (2018) Great tits and the city: distribution of genomic diversity and gene - environment associations along an urbanization gradient. Evolutionary Applications 11(5), 593–613.

PlayàMontmany, N. & Tattersall, G. (2021) Spot size, distance, and emissivity errors in field applications of infrared thermography. Methods in Ecology and Evolution Accepted.

Powers, D., Tobalske, B., Wilson, J., Wood, H. & Corder, K. (2015) Heat dissipation during hovering and forward flight in hummingbirds. Royal Society Open Science 2(12), 150598.

Prasher, S., Thompson, M., Evans, J., El-Nachef, M., Bonier, F. & Morand-Ferron, J. (2019) Innovative consumers: ecological, behavioral, and physiological predictors of responses to novel food. Behavioral Ecology 30(5), 1216–1225.

R Core Team (2019) R: A language and environment for statistical computing. R Foundation for Statistical Computing, Vienna, Austria.

Radchuk, V., Reed, T., Teplitsky, C., Van De Pol, M., Charmantier, A., Hassall, C., Adamík, P., Adriaensen, F., Ahola, M., Arcese, P. & Avilés, J. (2019) Adaptive responses of animals to climate change are most likely insu?cient. Nature Communications 10(1), 1–14.

Rao, F., Zhang, L., Wessel, J., Zhang, K., Wen, G., Kennedy, B., Rana, B., Das, M., Rodriguez-Flores, J., Smith, D. & Cadman, P. (2008) Adrenergic polymorphism and the human stress response. Annals of the New York Academy of Science 1148(1), 282.

Ratcliffe, L., Mennill, D. & Schubert, K. (2007) Social dominance and fitness in black-capped chickadees. Ecology and behavior of chickadees and titmice: an integrated approach, pp. 131–147, Oxford University Press, Oxford, Uk.

Rensel, M. & Schoech, S. (2011) Repeatability of baseline and stress-induced corticosterone levels across early life stages in the Florida scrub-jay (aphelocoma coerulescens). Hormones and Behavior 59(4), 497–502.

Rich, E. & Romero, L. (2005) Exposure to chronic stress downregulates corticosterone responses to acute stressors. American Journal of Physiology - Regulatory, Integrative and Comparative Physiology 288(1), R1628–R1636.

Richards, S. (1971) The significance of changes in the temperature of the skin and body core of the chicken in the regulation of heat loss. Journal of Physiology 216(1), 1–10.

Robb, G., McDonald, R., Chamberlain, D. & Bearhop, S. (2008) Food for thought: supplementary feeding as a driver of ecological change in avian populations. Frontiers in Ecology and the Environment 6(9), 476–484.

Robertson, J., Mastromonaco, G. & Burness, G. (2020a) Evidence that stress-induced changes in surface temperature serve a thermoregulatory function. Journal of Experimental Biology 223(4), jeb213421.

Robertson, J., Mastromonaco, G. & Burness, G. (2020b) Social hierarchy reveals thermoregulatory trade-o?s in response to repeated stressors. Journal of Experimental Biology 223(21), jeb229047.

Romero, L., Dickens, M. & Cyr, N. (2009) The reactive scope model - a new model integrating homeostasis, allostasis, and stress. Hormones and Behavior 55(3), 375–389.

Rouder, J., Morey, R., Speckman, P. & Province, J. (2012) Default bayes factors for ANOVA designs. Journal of Mathematical Psychology 56(5), 356–374.

Rutschmann, A.D., Ronce, O. & Chuine, I. (2015) Phenological plasticity will not help all species adapt to climate change. Global Change Biology 21(8), 3062–3073.

Schindelin, J., Arganda-Carreras, I., Frise, E., Kaynig, V., Longair, M., Pietzsch, T., Preibisch, S., Rueden, C., Saalfeld, S., Schmid, B. & Tinevez, J. (2012) Fiji: an open-source platform for biological-image analysis. Nature Methods 9(7), 676–682.

Sepp, T., McGraw, K., Kaasik, A. & Giraudeau, M. (2018) A review of urban impacts on avian lifehistory evolution: Does city living lead to slower pace of life? Global Change Biology 24(4), 1452–1469.

Seto, K., Güneralp, G. & Hutyra, L. (2012) Global forecasts of urban expansion to 2030 and direct impacts on biodiversity and carbon pools. Proceedings of the National Academy of Science 109(40), 16083–16088.

Seutin, G., White, B. & Boag, P. (1991) Preservation of avian blood and tissue samples for dna analyses. Canadian Journal of Zoology 9(1), 82–90.

Smit, B., Whitfield, M., Talbot, W., Gerson, A., McKechnie, A. & Wolf, B. (2018) Avian thermoregulation in the heat: phylogenetic variation among avian orders in evaporative cooling capacity and heat tolerance. Journal of Experimental Biology 221(6), jeb174870.

Srikanth, K., Kumar, H., Park, W., Byun, M., Lim, D., Kemp, S., Te Pas, M., Kim, J. & Park, J. (2019) Cardiac and skeletal muscle transcriptome response to heat stress in kenyan chicken ecotypes adapted to low and high altitudes reveal di?erences in thermal tolerance and stress response. Frontiers in Genetics 10(1), 993.

Tattersall, G. (2016) Infrared thermography: A non-invasive window into thermal physiology. Comparative Biochemistry and Physiology A: Molecular & Integrative Physiology 202(1), 78–98.

Tattersall, G. (2019) Thermimage: Thermal image analysis. R package version 4.0.1.

Thomas, C. (2017) Inheritors of the Earth: how nature is thriving in an age of extinction. PublicAffairs, New York, USA.

Thompson, M., Evans, J., Parsons, S. & Morand-Ferron, J. (2018) Urbanization and individual di?erences in exploration and plasticity. Behavioral Ecology 29(6), 1415–1425.

United Nations, Department of Economic and Social Affairs, Population Division (2019) World urbanization prospects: the 2018 revision (ST/ESA/SER.A/420). United Nations, New York, USA.

Van Homrigh, A., Higgie, M., McGuigan, K. & Blows, M. (2007) The depletion of genetic variance by sexual selection. Current Biology 17(6), 528–532.

Vasseur, D., DeLong, J., Gilbert, B., Greig, H., Harley, C., McCann, K., Savage, V., Tunney, T. & O’Connor, M. (2014) Increased temperature variation poses a greater risk to species than climate warming. Proceedings of the Royal Society B: Biological Sciences 281(1779), 20132612.

Vincent, J., Cardinaud, B. & Vernier, P. (1998) Evolution of monoamine receptors and the origin of motivational and emotional systems in vertebrates. Bulletin de L’Academie Nationale de Medecine 182(7), 1505–14.

Vincze, E., Seress, G., Lagisz, M., Nakagawa, S., Dingemanse, N. & Sprau, P. (2017) Does urbanization affect predation of bird nests? a meta-analysis. Frontiers in Ecology and Evolution 5(1), 29.

Visser, E., Van Reenen, C., Van der Werf, J., Schilder, M., Knaap, J., Barneveld, A. & Blokhuis, H. (2002) Heart rate and heart rate variability during a novel object test and a handling test in young horses. Physiology and Behavior 76(2), 289–296.

Von Borell, E., Langbein, J., Després, G., Hansen, S., Leterrier, C., Marchant-Forde, J., Marchant-Forde, R., Minero, M., Mohr, E., Prunier, A. & Valance, D. (2007) Heart rate variability as a measure of autonomic regulation of cardiac activity for assessing stress and welfare in farm animals = a review. Physiology and Behavior 92(3), 293–316.

Wagenmakers, E., Lodewyckx, T., Kuriyal, H. & Grasman, R. (2010) Bayesian hypothesis testing for psychologists: A tutorial on the Savage-Dickey method. Cognitive Psychology 60(3), 158–189.

Wan, J., Wang, C., Qu, H., Liu, R. & Zhang, Z. (2018) Vulnerability of forest vegetation to anthropogenic climate change in china. Science of the Total Environment 621(1), 1633–1641.

Watson, H., Videvall, E., Andersson, M. & Isaksson, C. (2017) Transcriptome analysis of a wild bird reveals physiological responses to the urban environment. Scientific Reports 7(1), 44180.

Weise, C. & Meyer, J. (1979) Juvenile dispersal and development of site-fidelity in the black-capped chickadee. The Auk 96(1), 40–55.

Wickham, H. (2016) ggplot2: elegant graphics for data analysis. Springer.

Winder, L., White, S., Nord, A., B, B.H. & McCafferty, D. (2020) Body surface temperature responses to food restriction in wild and captive great tits. Journal of Experimental Biology 223(8), jeb220046.

Wolak, M., Fairbairn, D. & Paulsen, Y. (2012) Guidelines for estimating repeatability. Methods in Ecology and Evolution 3(1), 129–137.

Wood, S. (2003) Thin plate regression splines. Journal of the Royal Statistical Society B 65(1), 95–114.

Yamamoto, K. & Vernier, P. (2011) The evolution of dopamine systems in chordates. Frontiers and Neuroanatomy 5(1), 21.

Yokoi, Y. (1966) Effect of ambient temperature upon emotional hyperthermia and hypothermia in rabbits. Journal of Applied Physiology 21(6), 1795–1798.

Zhuang, Z., Chen, S., Chen, C., Lin, E. & Huang, S. (2019) Genome-wide association study on the body temperature changes of a broiler-type strain Taiwan country chickens under acute heat stress. Journal of Thermal Biology 82(1), 33–42.

