## Supporting Information for "Thermal flexibility is a repeatable mechanism to cope with environmental stressors in a passerine bird"

#### 1 | Methods

##### 1.1 | Quantifying and comparing degree of urbanisation

To validate that our selected "urban" and "rural" sample locations were indeed representative of true urban and true rural ecotypes respectively, we used a similar approach to Thompson *et al.* (2018) with slight modifications to permit replication with free and open-source software (here, R statistical software; R Core Team 2019). Specifically, we first collected spatial data pertaining to broad land cover classification at each of our six capture locations, with capture locations representing plots of 0.05° longitude × 0.05° latitude (approximately 4.0 km by 5.5 km), centred on a given trapping location. Land cover classifications included in this study were: (1) water, (2) streets, (3) buildings, (4) crop-land, and (5) forests, and data for each classification type were obtained from Statistics Canada (2011, 2016, 2019), Agriculture and Agri-Food Canada (2015), and Ontario Ministry of Natural Resources and Forestry (2018) respectively. Remaining unclassified land was categorised as "bare ground" (including tarmack, lawn, and bare rock). Unfortunately, data pertaining to land-coverage by buildings was available for our urban capture locations alone. To correct for this problem, we collected satellite images (10 m resolution; Copernicus Sentinel-2, 2020) for the chosen range-limits of all capture locations using the R package "getSpatialData" (`https://jakob.schwalb-willmann.de/getSpatialData/`), then used these images and available building coverage data to construct a neural network algorithm to classify building presence or absence at each satellite image pixel (representing a 10 m × 10 m square), according to imaged light spectra. Here, our neural network algorithm was constructed using the R package "caret" (`https://cran.r-project.`

org/web/packages/caret/), using 10-fold cross-validation, with grid-search hyper-parameter tuning (size range = 5-15; decay range = 0.001-0.100). Importantly, to ensure that our algorithm was not solely constructed from theorised "urban" data, and thus was not biased by other surrounding land-coverage types that are typical of urban areas (e.g. lawns and tarmack), we manually determined the spatial locations of all identifiable buildings within one, randomly selected, rural capture location (Erin; 43.7617°N, 80.1529°W; range limits described above) using a freely available, online, geospatial polygon selection tool (http://apps.headwallphotonics.com/), then included these data within our total, building coverage data-set. Next, we partitioned our building coverage data-set into training data (75% of all data) that was used for algorithm construction, and testing data (25% of all data) that was used for algorithm validation (refer to Results below). Finally, satellite image data for all capture locations were loaded into our neural network algorithm and resultant predictions of building coverage were used in conjunction with water, street, crop-land, forest, and bare ground categorisations for subsequent analyses.

To generate a univariate metric for degree of urbanisation, we again followed methods described by Thompson *et al.* (2018). To do so, we summed the number of 10 m × 10 m squares (representing the size of a satellite image pixel) within each capture location that pertained to a given land coverage type (classifications described above). We then sought to load these count data into a principle component analysis ("PCA") and subsequently extract values from the principle component ("PC") that best represented a degree of urbanisation (e.g. high building and road coverage, and low water, crop-land, and wooded area coverage), for each capture location. Given the small number of urban and rural capture locations used in this study (n = 3 locations per theorised ecotype), however, we could not confidently estimate variance of, and covariance between, count numbers for land coverage categorisations across capture locations. To therefore increase our sample size for PCA construction, we randomly selected two expectedly urban (cities of Hamilton and Kitchener; 43.1339°N, 79.5210°W, and 43.2737°N, 80.3155°W respectively) and two expectedly rural locations (the township of Gowanstown and the Luther Marsh wildlife area; 42.4543°N, 80.5118°W and 43.5681°N, 80.2511°W, respectively) within southern Ontario, Canada (maximum distance between random sample locations and the closest true capture location = 57.15 km), then classified land coverage within each sample location using both available data-sets and satellite imagery described above. Finally, the number of 10 m x 10 m squares pertaining to each land coverage classification were again summed, and all sums were subsequently loaded into a scaled and centred PCA in R. Similar to Thompson *et al.* (2018) principle component one ("PC1") best represented the degree of urbanisation within a capture location (Fig. C2) and explained 58.93% of count variance. We therefore extracted values of PC1 (henceforth, "degree of urbanisation") for each of our six true capture locations, then scaled these values between 0 and 100 (0 = highly rural; 100 = highly urban) to aid in visual interpretation.

To statistically test whether average degree of urbanisation was larger in our expectedly urban capture locations than that in our expectedly rural capture locations, we calculated Bayes factors (K) for a non-linear hypothesis test using the Savage-Dickey method (Wagenmakers *et al.*, 2010). Bayes factors provide an indication of the relative support for one hypothesis over another (here, that the degree of urbanisation differed between *a priori* ecotype classifications, or did not), and were therefore considered a useful statistic for validating our *a priori* assumptions of ecotype classification. Prior distributions for the degree of urbanisation at each capture location were assumed to be normal with means of 50 and standard deviations (s.d.) of 2.

#### 1.2 | Testing the effects of resource acquisition on repeatability of stress-induced thermal responses

Recent research has shown that stress-induced changes in surface temperature may represent true trade-offs between the physiological stress response and thermoregulation (Oka, 2018; Robertson *et al.*, 2020). According to this hypothesis, both the presence and magnitude of stress-induced thermal responses are thought to negatively reflect resource availability, with individuals experiencing a limitation of resources displaying more pronounced thermal responses to stress exposure than non-resource limited conspecifics (Robertson *et al.*, 2020). As such, consistent differences in both acute and chronic stress-induced thermal responses that we report in our study (see "Results" section) may merely reflect consistent differences in resource availability (an "environmental" effect) and not differences in genetic or epigenetic architecture among individuals.

To test whether relative access to resources could indeed explain consistent differences in stress-induced thermal responses in our sample population, we first re-estimated the repeatabilities of both the acute and chronic stress-induced thermal responses among black-capped chickadees (i.e. repeatabilities of stress-induced changes in surface temperature and heat-loss; see "Statistical Analysis" section of the main text), while replacing the true identities of individual chickadees with those of other chickadees that share a similar rate of resource acquisition. These new repeatability estimates (or "fixed resource repeatability estimates"), therefore, should reflect those explained by differences in the relative access of resource among individuals alone, and not differences in genetic or epigenetic identity. In our sample population, access to resources (here, proxied by feeding rate) is known to vary according to social status, with socially dominant individuals feeding more frequently than socially subordinate individuals

(Robertson *et al.*, 2020). Thus, identities of chickadees were randomly scrambled according to known social status (see Robertson *et al.* 2020) and according to treatment order (see "Statistical Analysis" section of the main text). Next, we compared our new, fixed resource repeatability estimates with our previous "true" repeatability estimates (i.e. where individual identity remained unscrambled; again, see "Statistical Analysis" section of the main text) using four, one-way non-linear hypothesis tests in the R package "brms" (Bürkner, 2017). Priors for our non-linear hypothesis tests were beta distributed with peaks at 0 ($\alpha$ = 1, and $\beta$ = 4), and Bayes factors for our hypothesis tests were calculated using the Savage-Dickey density ratio method Wagenmakers *et al.* (2010).

For our non-linear hypothesis tests, a significant increase in our true repeatability estimates relative to our fixed resource repeatability estimates would indicate that consistent variation in stress-induced thermal responses among individuals cannot be explained by variations in resource access alone. By contrast, a statistical similarity (or non-significant difference) between our true and fixed resource repeatability estimates would indicate that consistent variation in stress-induced thermal responses among individuals may be largely explained by variations in resource access among individuals.

### 2 | Results

Estimates are reported alongside 95% credible intervals in carets.

#### 2.1 | Neural network algorithm validation

Accuracy of our algorithm when detecting building coverage using satellite image data was 90.09% [0.899, 0.903]. Cohen's $\kappa$ was estimated as 44.42%, suggesting that precision of building detection by our algorithm was moderately high. Finally, sensitivity of our algorithm was moderate (39.06%), while specificity was high (97.33%).

#### 2.2 | Comparison of degree of urbanisation

The degree of urbanisation ranged from 43.84 to 100 in our expectedly urban capture locations, and from 0 to 7.38 in our expectedly rural capture locations. Our non-linear hypothesis test strongly supported a positive difference between degree of urbanisation at our expectedly urban capture locations and degree of urbanisation at our expectedly rural capture locations (K > 100; $\beta$ = 67.98 [42.09, 96.43]; Figures S3-S4).

#### 2.3 | Variations in access to resources are insufficient to explain consistent variations in stress-induced thermal responses

Across acute time periods ($\leq$ one hour), repeatability estimates derived from our true models (i.e. "true repeatability estimates") were significantly larger than those drawn from data where individual identities were randomly swapped among birds with similar rates of resource acquisition (i.e. "fixed resource repeatability estimates). Specifically, the repeatabilies of stress-induced changes in surface temperature and heat-loss estimated from our true data were 0.14 [0.03,0.32] and 0.11 [0.02, 0.27] respectively, while those estimated from randomly generated data were 0.01 [0.00, 0.03] and 0.01 [0.00, 0.03] respectively (Figure S11). Bayes factors (representing the evidence that true repeatability estimates outweigh fixed resource repeatability estimates) were > 100 and 96.3 for comparisons of stress-induced changes in surface temperature and heat-loss respectively. Across chronic time periods (> one hour and $\leq$ 30 days), similar trends were observed. Namely, the repeatability of stress-induced changes in surface temperature and heat-loss estimated from our true data (surface temperature: R = 0.61 [0.35, 0.81]; heat-loss: R = 0.67 [0.44, 0.84]; Figure S12a) significant outweighed those estimated from our randomly generated data (surface temperature: R = 0.03 [0.00, 0.09]; heat-loss: R = 0.03 [0.00, 0.10]; Figure S12b). Together, these results suggest that variations in rates of resource acquisition among individuals were insufficient to explain consistent variations in stress-induced thermal responses at both acute and chronic time periods ($K_{surface\ temperature}$ > 100; $K_{heat\text{-}loss}$ > 100).

### 3 | Figures

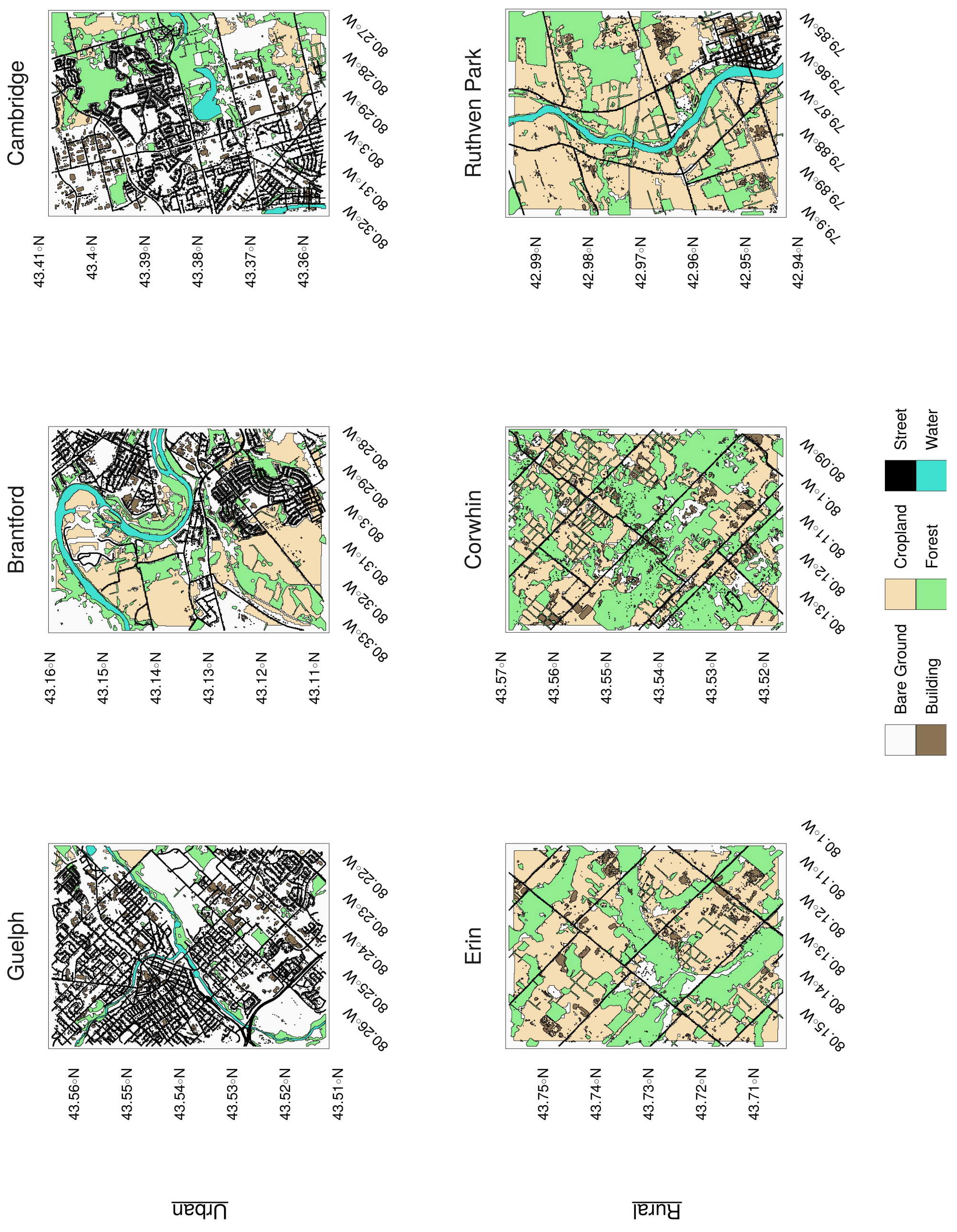
Cambridge
Brantford
Guelph
Ruthven Park
Corwhin
Erin
Urban
Rural
Bare Ground
Building
Cropland
Forest
Street
Water

FIGURE S1 **Locations in Ontario, Canada, at which black-capped chickadees used in experimentation were captured.** Maps represent precise trapping locations (centre of a give map) ± 0.025° latitude and 0.025° longitude. Data pertaining to land coverage by cropland, streets, water and forest were obtained from Statistics Canada (2011, 2016, 2019), Agriculture and Agri-Food Canada (2015), and the Ontario Ministry of Natural Resources and Forestry (2018) respectively. Land coverage pertaining to buildings was estimated from a semi-automated, neural network algorithm (accuracy = 90%).

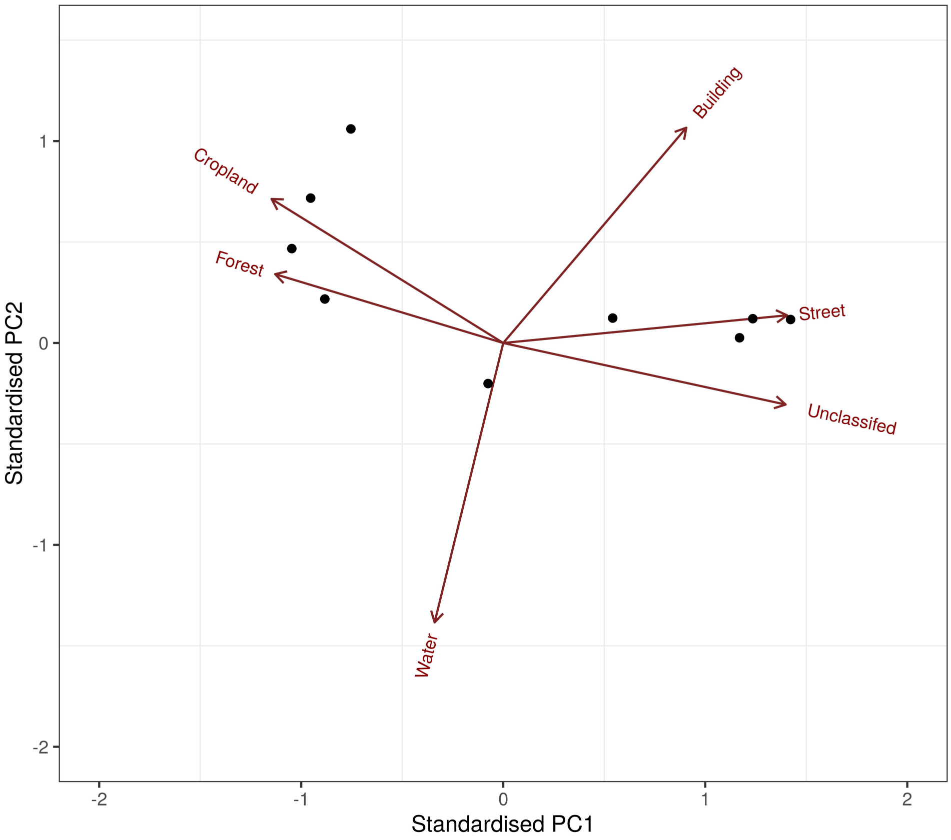


FIGURE S2 **Biplot of principle component analysis with relative quantity of land coverage categories at given sample locations (n = 10) as loading variables.** Principle component analysis is scaled and centred, and sample locations include true black-capped chickadee capture locations (n = 6) and randomly selected, nearby areas of equivalent size (n = 4).

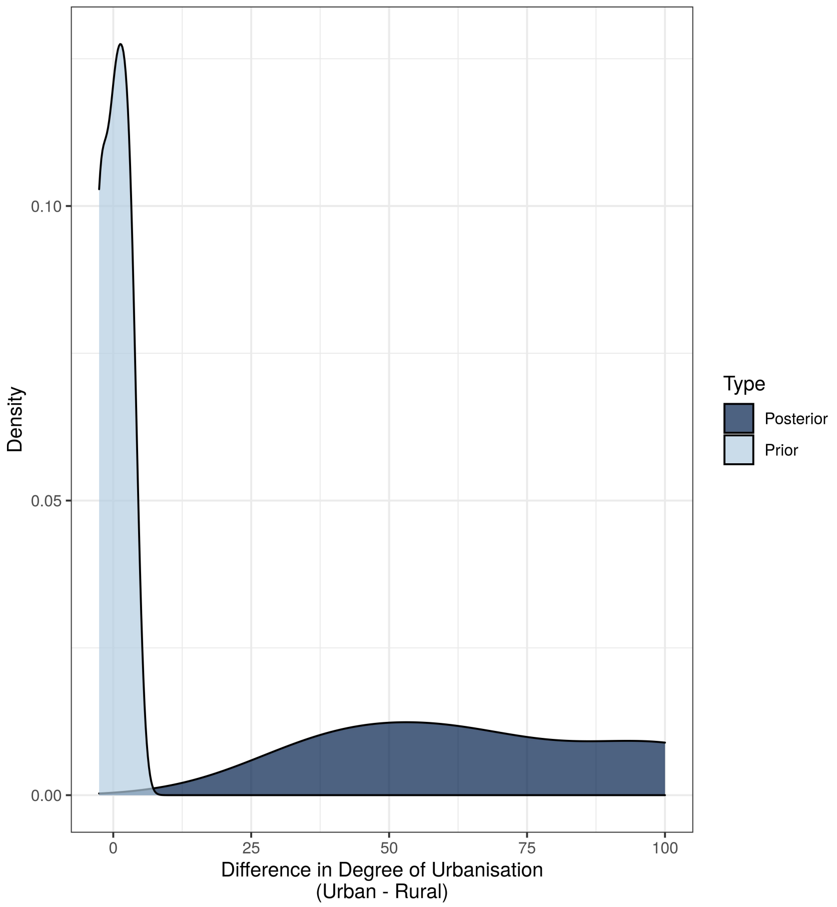


FIGURE S3 **Distribution of differences between degrees of urbanisation at expectedly urban and expectedly rural capture locations (n = 6).** Posterior differences represent true paired sample differences, while prior differences are estimated from random, normal distributions per capture ecotype (i.e. urban or rural; mean = 50; s.d. = 2).

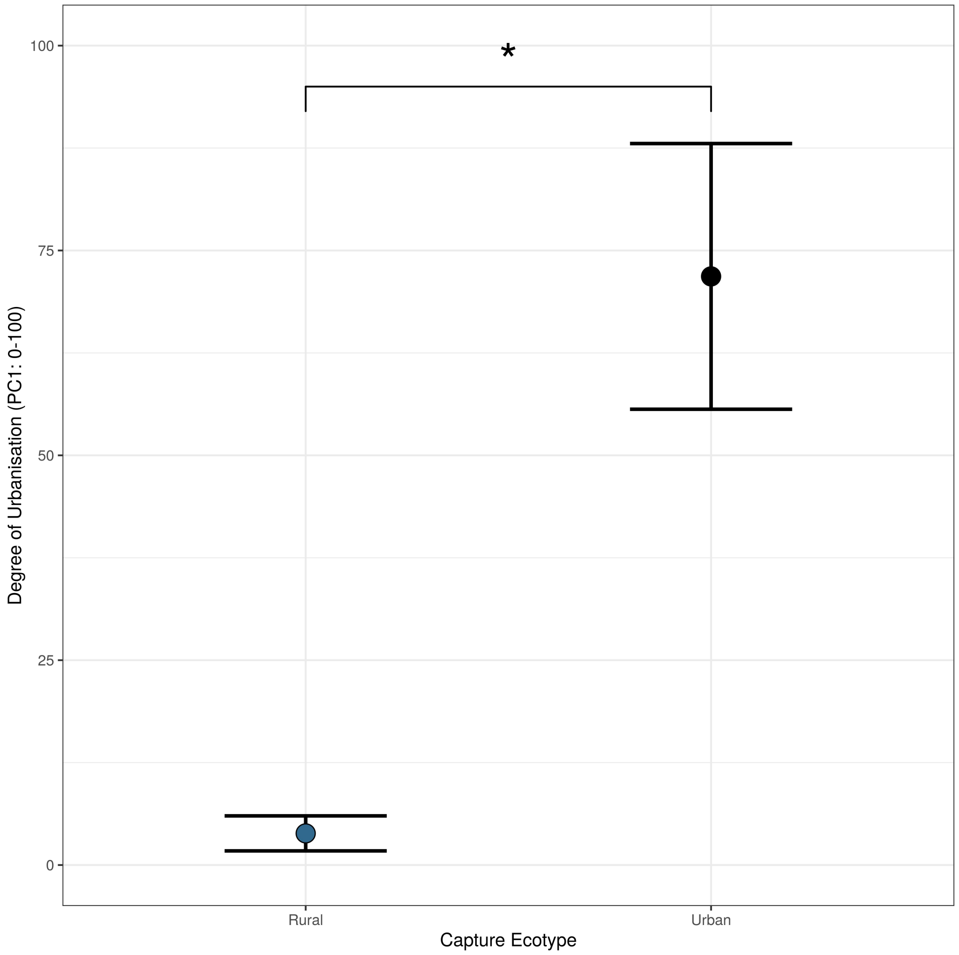


FIGURE S4 **Mean degree of urbanisation at expectedly urban (n = 3) and expectedly rural (n = 3) capture locations.** Degree of urbanisation was obtained from a principle component analysis with total area of water, forest, streets, crop-land, bare ground, and buildings within a capture location as loading variables. A low degree of urbanisation represents a highly rural ecotype (i.e. low densities of roads, buildings, and bare earth), while a high degree of urbanisation represents a highly urban ecotype (i.e. high densities of roads, buildings, and bare earth). Whiskers represent means $\pm$ standard errors of means.

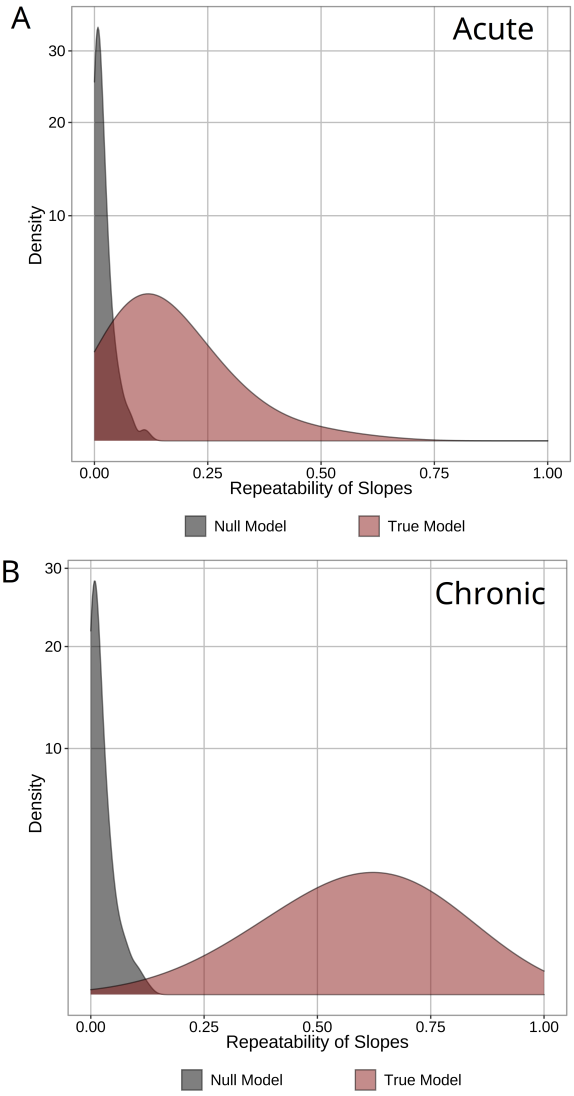


FIGURE S5 **Repeatability of acute and chronic changes in eye region temperature ($T_s$) following stress exposure in black-capped chickadees (n = 19).** Panels **A** and **B** display distribution of true and null repeatability estimates for acute and chronic thermal responses to stress exposure respectively. Distributions represent density of repeatability values estimated from posterior distributions of hierarchical generalised additive mixed effects model (GAMM). Thermal responses to stress exposure represent those observed at the eye region of chickadees, using infra-red thermography across 60 days of observation.

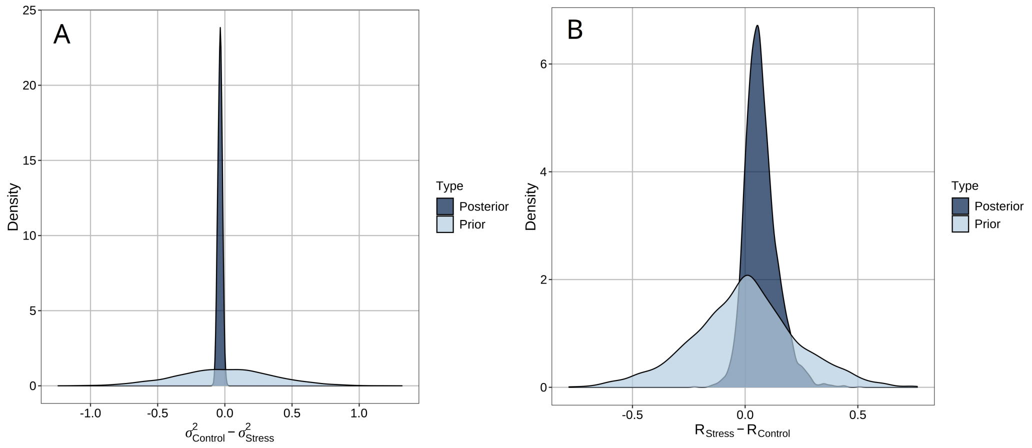


FIGURE S6 **Distribution of differences between unexplained variability and repeatability of eye region temperature ($T_s$) measurements derived from black-capped chickadees (n = 19) in control and acute stress-exposed treatments.** Differences ($n_{Acute}$ = 3600; $n_{Chronic}$ = 3600) are drawn from posterior and prior distributions from Bayesian generalised additive mixed effects models (GAMMs). **A |** Distribution of differences between unexplained variability (residual error) in $T_s$ during control treatments and unexplained variability in $T_s$ during stress exposure treatments. **B |** Distribution of differences between repeatability of $T_s$ across time in stress exposure treatments, and repeatability of $T_s$ across time in control treatments.

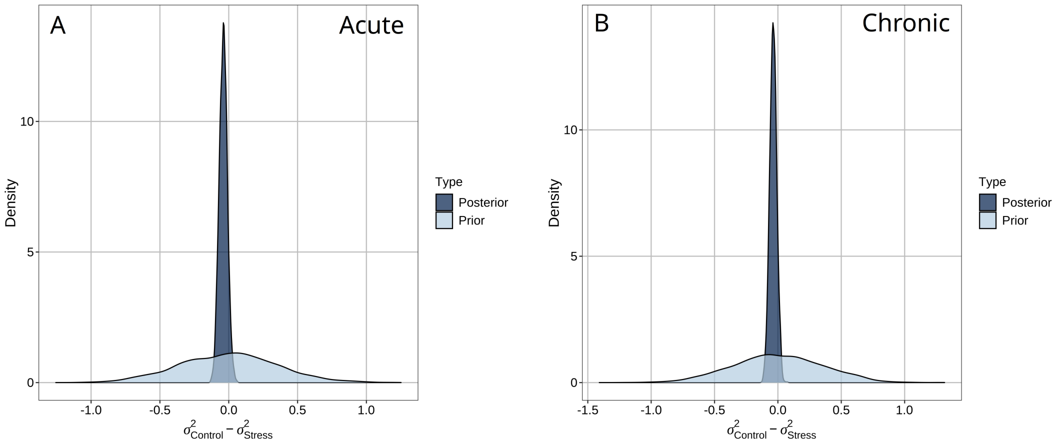


FIGURE S7 **Distribution of differences between unexplained variability and repeatability of heat transfer at the eye region ($q_{Tot}$) among black-capped chickadees (n = 19) in chronic and acute stress-exposed treatments.** Differences ($n_{Acute}$ = 3600; $n_{Chronic}$ = 3600) are drawn from posterior and prior distributions from Bayesian generalised additive mixed effects models (GAMMs). **A |** Distribution of differences between unexplained variability (residual error) in $q_{Tot}$ during control treatments and unexplained variability in $q_{Tot}$ during stress exposure treatments. **B |** Distribution of differences between repeatability of $q_{Tot}$ across time in stress exposure treatments, and repeatability of $q_{Tot}$ across time in control treatments.

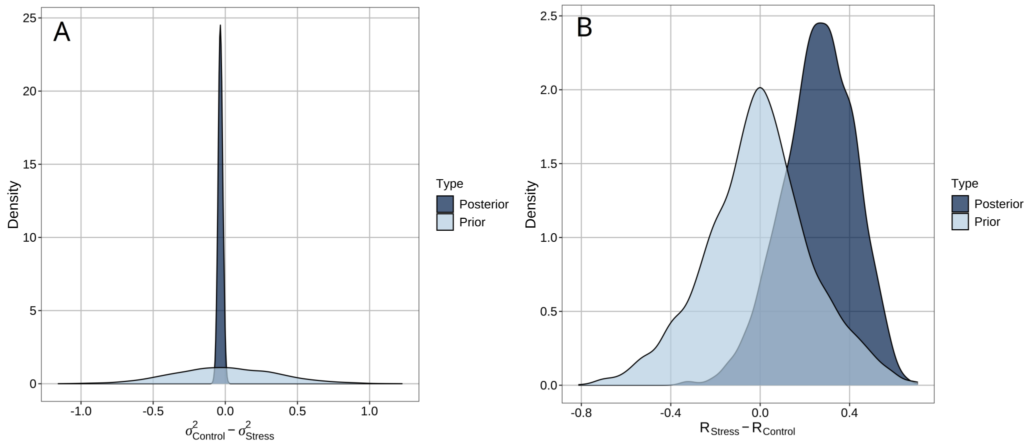


FIGURE S8 **Distribution of differences between unexplained variability and repeatability of eye region temperature ($T_s$) measurements derived from black-capped chickadees (n = 19) in control and chronic stress-exposed treatments.** Differences ($n_{Acute}$ = 3600; $n_{Chronic}$ = 3600) are drawn from posterior and prior distributions from Bayesian generalised additive mixed effects models (GAMMs). **A |** Distribution of differences between unexplained variability (residual error) in $T_s$ during control treatments and unexplained variability in $T_s$ during stress exposure treatments. **B |** Distribution of differences between repeatability of $T_s$ across ambient temperature in stress exposure treatments, and repeatability of $T_s$ across ambient temperature in control treatments.

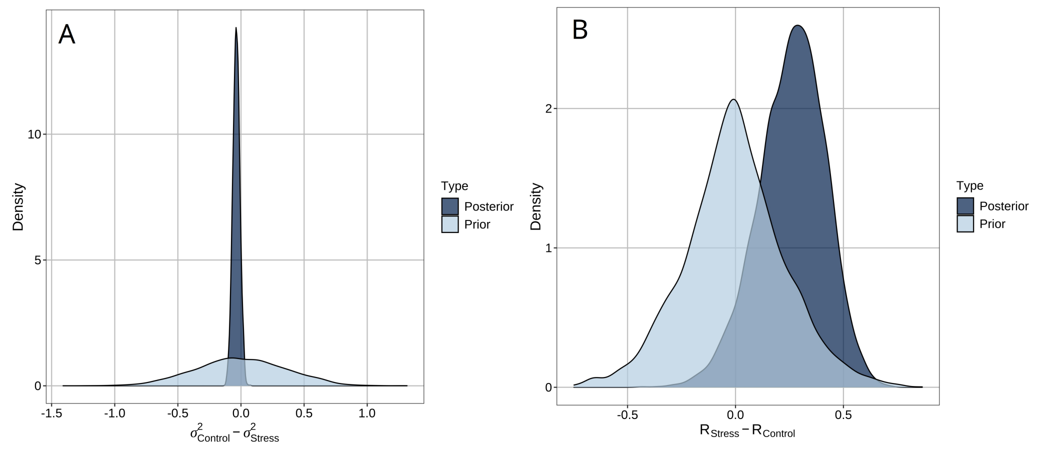


FIGURE S9 **Distribution of differences between unexplained variability and repeatability of heat transfer at the eye region ($q_{Tot}$) among black-capped chickadees (n = 19) in chronic and chronic stress-exposed treatments.** Differences ($n_{Acute}$ = 3600; $n_{Chronic}$ = 3600) are drawn from posterior and prior distributions from Bayesian generalised additive mixed effects models (GAMMs). **A |** Distribution of differences between unexplained variability (residual error) in $q_{Tot}$ during control treatments and unexplained variability in $q_{Tot}$ during stress exposure treatments. **B |** Distribution of differences between repeatability of $q_{Tot}$ across ambient temperature in stress exposure treatments, and repeatability of $q_{Tot}$ across ambient temperature in control treatments.

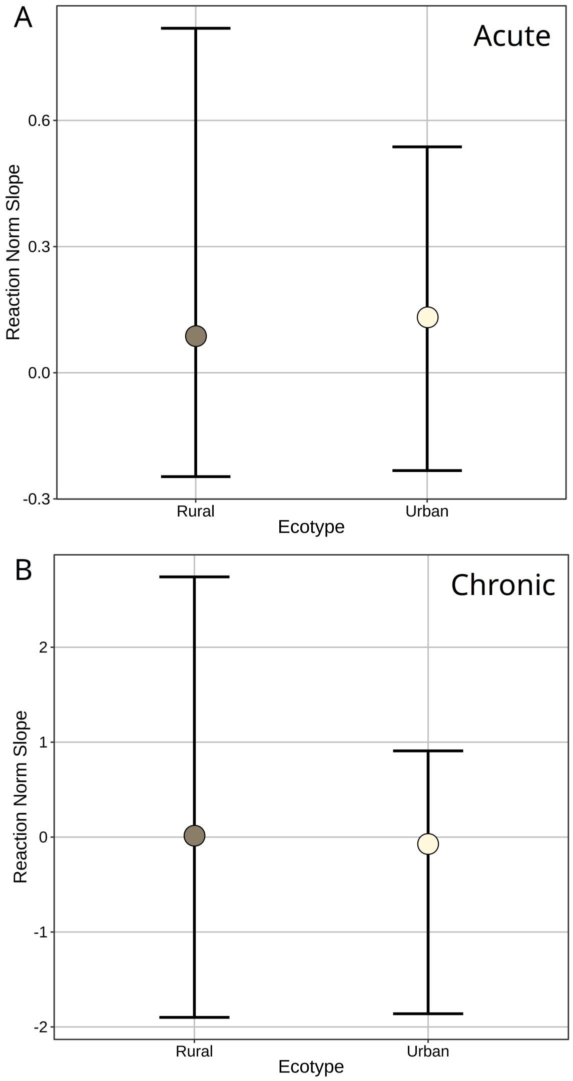


FIGURE S10 **Average effects of stress exposure on surface temperature ($T_s$; reaction norm slopes) at the eye region of black-capped chickadees (n = 19) captured from urban and rural ecotypes (n = 9 urban, n = 10 rural). A |** Average slopes of acute reaction norms across individuals captured at each ecotype. Reaction norm slopes represent the slopes of the linear interaction between treatment type and time post stress exposure (s) per individual. **B |** Average slopes of chronic reaction norms across individuals captured from each ecotype. Here, reaction norm slopes represent those of linear interactions between treatment type and ambient temperature (°C) per individual. Error bars represent 95% credible intervals around mean estimates. All reaction norm slopes were derived from Bayesian generalised additive mixed effects models (GAMMs).

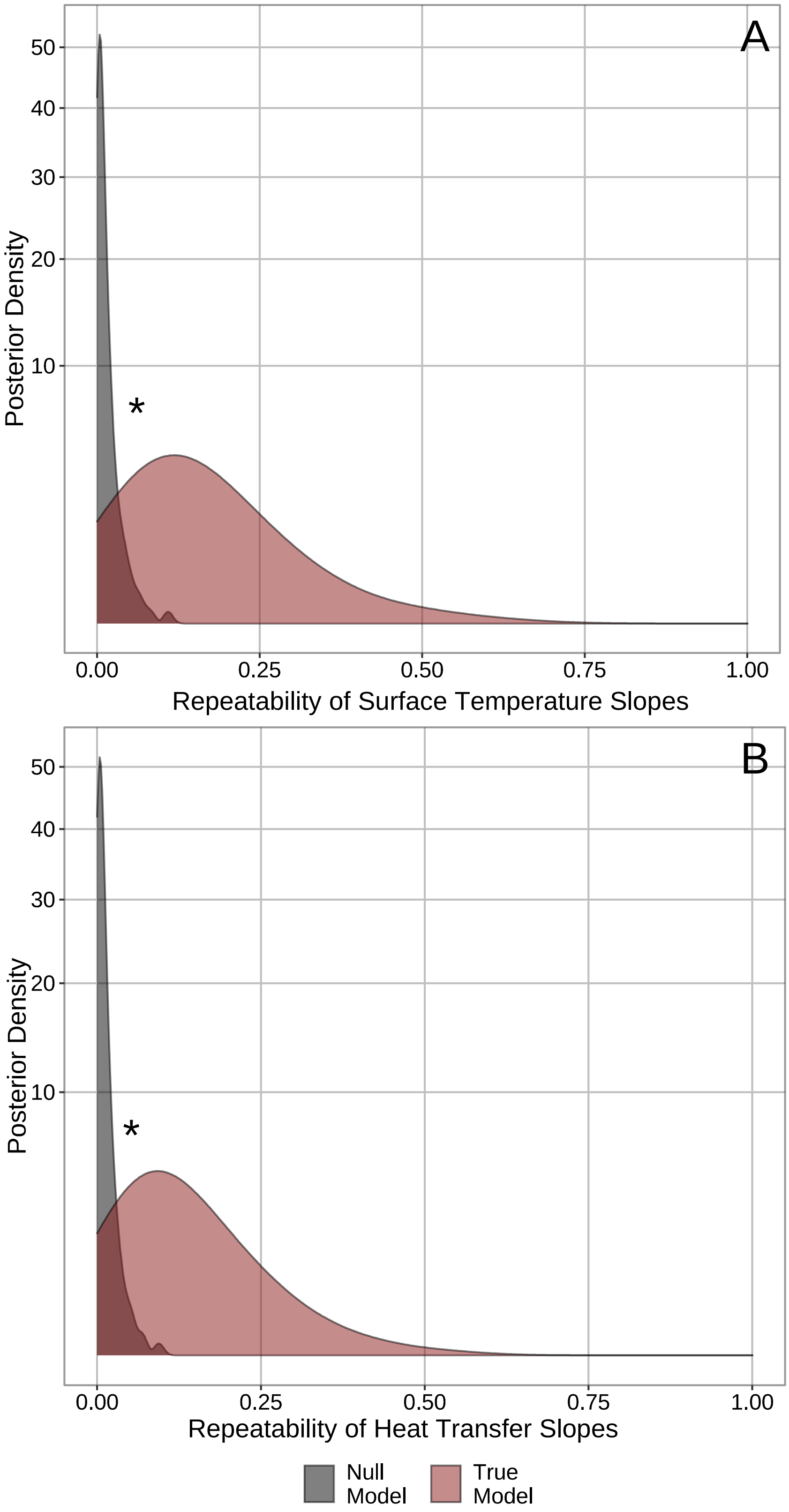
A
50
40
30
20
10
Posterior Density
*
0.00
0.25
0.50
0.75
1.00
Repeatability of Surface Temperature Slopes
B
50
40
30
20
10
Posterior Density
*
0.00
0.25
0.50
0.75
1.00
Repeatability of Heat Transfer Slopes
Null Model
True Model

FIGURE S11 **Repeatability of stress-induced thermal responses across acute time-periods ($\leq$ 1 hour) as estimated from real data ("true model"), and randomly generated data that controls for variations in resource access among individuals ("null model").** Panels **A** and **B** represent distribution of repeatability values estimated for stress-induced changes in surface temperature and stress-induced changes in dry heat-loss, respectively. True model distributions (red) represent those of drawn from models where identity of individuals was correctly identified. Null model distributions (grey) represent those drawn from models where identity of individuals was randomly shuffled among those with the same dominance status (and thus, rate of resource acquisition; Robertson et al, 2020b). A positive difference between true and null distributions (indicated by an asterisk, "*") implies that repeatability values from true models cannot be explained by biases in experimental methods (captured in null models) and are considered significant. Distributions are estimated from posteriors of Bayesian generalised additive mixed effects models (GAMM). Thermal responses to stress exposure represent those observed at the eye region of black-capped chickadees (n = 19), using infra-red thermography across 60 days of observation.

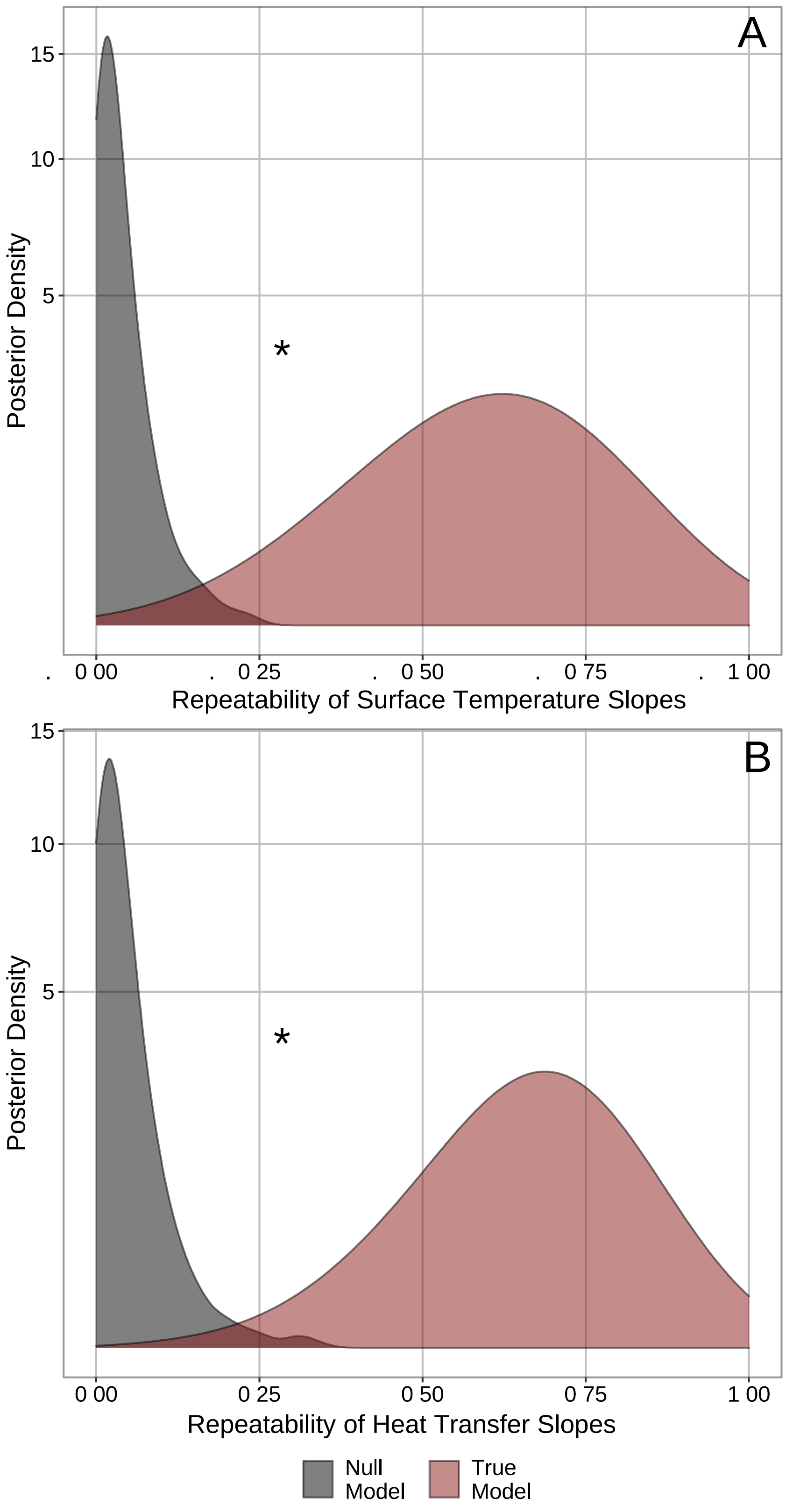
A
15
10
5
Posterior Density
*
. 0 00 . 0 25 . 0 50 . 0 75 . 1 00
Repeatability of Surface Temperature Slopes
B
15
10
5
Posterior Density
*
0 00 0 25 0 50 0 75 1 00
Repeatability of Heat Transfer Slopes
Null Model
True Model

FIGURE S12 **Repeatability of stress-induced thermal responses across chronic time-periods (> 1 hour and $\leq$ 30 days) as estimated from real data ("true model"), and randomly generated data that controls for variations in resource access among individuals ("null model").** Panels **A** and **B** represent distribution of repeatability values estimated for stress-induced changes in surface temperature and stress-induced changes in dry heat-loss, respectively. True model distributions (red) represent those of drawn from models where identity of individuals was correctly identified. Null model distributions (grey) represent those drawn from models where identity of individuals was randomly shuffled among those with the same dominance status (and thus, rate of resource acquisition; Robertson et al, 2020b). A positive difference between true and null distributions (indicated by an asterisk, "*") implies that repeatability values from true models cannot be explained by biases in experimental methods (captured in null models) and are considered significant. Distributions are estimated from posteriors of Bayesian generalised additive mixed effects models (GAMM). Thermal responses to stress exposure represent those observed at the eye region of black-capped chickadees (n = 19), using infra-red thermography across 60 days of observation.

### References

Agriculture and Agri-Food Canada (2015) Land use 1990, 2000, and 2010 (LU1990, LU2000, LU2010). url: https://open.canada.ca/data/en/dataset/9e1efe92-e5a3-4f70-b313-68fb1283eadf .

Bürkner, P. (2017) brms: An r package for bayesian multilevel models using stan. *Journal of Statistical Software* **80(1)**, 1–28.

Oka, T. (2018) Stress-induced hyperthermia and hypothermia. *Handbook of Clinical Neurology* (ed. A. Romanovsy), vol. 157, pp. 599–621, Elsevier, Amsterdam, Netherlands.

Ontario Ministry of Natural Resources and Forestry (2018) Land information Ontario: wooded areas .

R Core Team (2019) *R: A language and environment for statistical computing*. R Foundation for Statistical Computing, Vienna, Austria.

Robertson, J., Mastromonaco, G. & Burness, G. (2020) Social hierarchy reveals thermoregulatory trade-offs in response to repeated stressors. *Journal of Experimental Biology* **223(21)**, jeb229047.

Statistics Canada (2011) *Boundary Files, 2011 Census*. Statistics Canada Catalogue no. 92.160-X.

Statistics Canada (2016) *Road network files, 2016 Census*. Statistics Canada Catalogue no. 92.500-G, url: https://www12.statcan.gc.ca/census-recensement/2011/geo/RNF-FRR/index-2011-eng.cfm?year=16.

Statistics Canada (2019) *Open database of buildings (database)*. url: https://www.statcan.gc.ca/eng/ lode/databases/odb.

Thompson, M., Evans, J., Parsons, S. & Morand-Ferron, J. (2018) Urbanization and individual differences in exploration and plasticity. *Behavioral Ecology* **29(6)**, 1415–1425.

Wagenmakers, E., Lodewyckx, T., Kuriyal, H. & Grasman, R. (2010) Bayesian hypothesis testing for psychologists: A tutorial on the Savage-Dickey method. *Cognitive Psychology* **60(3)**, 158–189.
